# Weak SLP-76-PLC-γ1 interaction in the LAT-nucleated multi-protein complex fine-tunes TCR signal strength to optimize T cell responsiveness

**DOI:** 10.1101/2025.09.11.675682

**Authors:** Hidehiro Yamane, Junya Wada, Elizabeth N. Stassenko, Natalie D. Convertino, Mariah E. Lee, Melanie S. Vacchio, Wenmei Li, Udumbara M. Rathnayake, Lakshmi Balagopalan, Raj Chari, Herbert Hagenau, Parirokh Awasthi, Dorian B. McGavern, Remy Bosselut, Lawrence E. Samelson

**Affiliations:** Laboratory of Cellular and Molecular Biology, Center for Cancer Research (CCR), National Cancer Institute (NCI), National Institutes of Health (NIH), Bethesda, MD, U.S.A; Laboratory of Integrative Cancer Immunology, CCR, NCI, NIH, Bethesda, MD, U.S.A; Genome Modification Core, Laboratory of Animal Sciences Program (LASP), Frederick National Laboratory for Cancer Research (FNLCR), NCI, NIH, Frederick, MD, U.S.A; Mouse Modeling and Cryopreservation, LASP, FNLCR, NCI, NIH, Frederick, MD, U.S.A; Viral Immunology & Intravital Imaging Section, Neuroimmunology & Neurovirology Division, National Institute of Neurological Disorders and Stroke, NIH, Bethesda, MD, U.S.A

## Abstract

Upon TCR engagement several protein tyrosine kinases are recruited and activated, and adapter proteins and enzymes are phosphorylated on tyrosine residues, leading to further events characterizing activated T cells. Phosphorylation of the LAT adapter protein enables binding of the enzyme PLC-γ1 and of a dimer of two additional adapter proteins Gads and SLP-76, forming a tetrameric structure. Within this heterotetramer there is a weak interaction between SLP-76 and PLC-γ1, and the relevant binding sites of SLP-76 and PLC-γ1 are highly conserved in vertebrates. To address the biological relevance of this weak interaction, we introduced a mutation in the SLP-76 that enhanced its affinity for PLC-γ1 and found that this mutation increased PLC-γ1 activity and altered thymocyte development and peripheral T cell responses due to enhanced TCR signal strength. The conserved weak SLP-76-PLC-γ1 interaction is critical for the controlled activation of PLC-γ1, thus fine-tuning TCR signal strength to optimize T cell-mediated immunity.

## Introduction

The engagement of the T cell antigen receptor (TCR) with antigenic peptide bound to an MHC-encoded protein is the initiating event in the activation of T cells. Recruitment of critical protein tyrosine kinases (PTKs) including ZAP-70 to the intracellular domains of the TCR and activation of the PTKs rapidly follow. These PTKs have multiple enzyme and adapter protein substrates. We have focused on the consequences of the ZAP-70-mediated tyrosine phosphorylation of the integral membrane adapter protein LAT^1^. The phosphorylation of this molecule on four of its intracellular tyrosine residues creates binding sites for additional adapter molecules and enzymes, leading to activation of multiple intracellular events. Phospholipase C-γ1 (PLC-γ1) is one such critical enzyme. It translocates from the cytoplasm to the plasma membrane to bind phosphorylated LAT upon T cell activation. Additionally, a dimer of two cytosolic adapters Gads and SLP-76 also binds to phospho-LAT and PLC-γ1, thus creating a tetrameric protein complex, LAT-Gads-SLP-76-PLC-γ1^2, 3^. Upon TCR-mediated activation multiple additional proteins including the adapters Nck and Vav and the enzyme ITK bind to phosphorylated SLP-76 at the LAT-nucleated tetrameric complex in order to phosphorylate and thus activate PLC-γ1 to target its substrate phosphatidylinositol 4,5-bisphosphate (PIP2) at the plasma membrane. Cleavage of this lipid results in the release of inositol trisphosphate (IP3), which mediates the elevation of intracellular calcium, and diacylglycerol which leads to the activation of PI-3 kinase and critical protein serine kinases^4^.

We have studied the formation of the tetrameric complex *in vitro* using recombinant and synthetic proteins, isothermal titration calorimetry (ITC) and analytical ultracentrifugation (AUC) to determine the thermodynamics of protein-protein interactions and tetramer formation^3,5^. The individual affinities of binding between phospho-LAT and Gads, Gads and SLP-76, and phospho-LAT and PLC-γ1 are in the sub-micromolar range, though the interaction of SLP-76 with PLC-γ1 is undetectable at room temperature and barely detected at 4°C with a K_d_ of 3.3 µM^5^. Using all four proteins we subsequently demonstrated the formation of the tetrameric complex by AUC and showed that the concentration range of tetramer formation is estimated to be 0.1-1 µM^3^. We observed that the favorable decrease in enthalpy generated during tetramer formation mostly compensates for the entropic penalty resulting from the spatially limited circular arrangement of the tetramer^3^. We suggested that the tetramer is in thermodynamic equilibrium between open and closed circular structures regulated by the SLP-76-PLC-γ1 interaction. In a second study we used liposome-based reconstitution to define the membrane recruitment and enzymatic activities of these proteins. Formation of the tetramer at the liposomal membrane enhances phosphoinositide hydrolysis by PLC-γ1 over that induced by PLC-γ1 alone^6^. These results led us to propose the model that the formation of a closed circular tetrameric structure mediated by the SLP-76-PLC-γ1 interaction induces a conformational change in PLC-γ1 that activates its enzymatic function.

Based on our previous findings, we hypothesized that the low affinity interaction between SLP-76 and PLC-γ1 in the setting of the tetramer might serve as a critical check point for T cell immune responses. In support of this hypothesis, the animo acid sequences of the interaction sites of SLP-76 and PLC-γ1 were highly conserved among jawed vertebrates. Therefore, we sought to determine the consequences of increasing the affinity of interaction between these two components by substituting the relevant region of SLP-76 with a short sequence of amino acids (aa) previously demonstrated to have a higher affinity for the PLC-γ1 SH3 domain^7^. We found that this higher affinity interaction increased the activity of PLC-γ1 *in vitro*. We then generated a mouse in which the higher affinity proline-rich region (PRR) of SLP-76 replaced the wild-type (WT) sequence specifically in T cells and observed impaired thymocyte development. Moreover, using TCR transgenic systems, we found that this mutation led to altered CD4^+^ and CD8^+^ T cell immune responses. We conclude that the low affinity interaction between SLP-76 and PLC-γ1 is an evolutionally conserved mechanism that underlies the fine-tuning of the strength of TCR-mediated signaling to optimize T cell development and T cell-mediated immunity.

## Results

### In vitro reconstitution and characterization of the SLP-76-PLC-γ1 interaction

In previous studies of intracellular signaling downstream of the TCR, we used *in vitro* reconstitution techniques to demonstrate and verify that four signaling proteins, PLC-γ1, LAT, Gads, and SLP-76, form a heterotetrameric protein complex through the following interactions: the SH2 domain of PLC-γ1 and pY132 of LAT, the SH2 domain of Gads and pY171 of LAT, the PRR_225-250_ of SLP-76 and the SH3 domain of Gads, and the PRR_185-200_ of SLP-76 and the SH3 domain of PLC-γ1^3, 5^ (Fig. 1a). Calorimetric data for the protein-protein interactions defining this tetramer are shown in Table 1. Of note is that the interaction between PLC-γ1 and SLP-76 is undetectable at room temperature and has a K_d_ of only 3345 nM at 4℃^3, 5^. To examine the significance of the interaction between SLP-76 and PLC-γ1 over evolution in more detail we compared the protein sequences of nineteen orthologs of SLP-76 and PLC-γ1 in jawed vertebrates, ranging from cartilaginous fish to placental mammals. These sequences were aligned by Protein BLAST (Fig. 1b and Extended Data Fig. 1a). The SH3 domain of PLC-γ1 binds to the PRR of SLP-76 (aa 180-215 from human sequence). In this PRR, two double prolines (PP) and an arginine-proline pair (RP) were perfectly conserved, which is consistent with the previous reports about the importance of the PXXP motif in this region^7, 8^ (Fig. 1b). As this region interacts with the SH3 domain of PLC-γ1 (aa 791-851 from human sequence), we also aligned the sequences of vertebrate PLC-γ1 SH3 domains interacting with SLP-76 and found near perfect conservation (Extended Data Fig. 1a). The sequence conservation of the relevant regions of the SLP-76 PRR and the PLC-γ1 SH3 domain suggest a crucial role of this protein-protein interaction during the cooperative activation of PLC-γ1 by tetramer formation upon TCR engagement^3, 5^.

**Figure 1:**
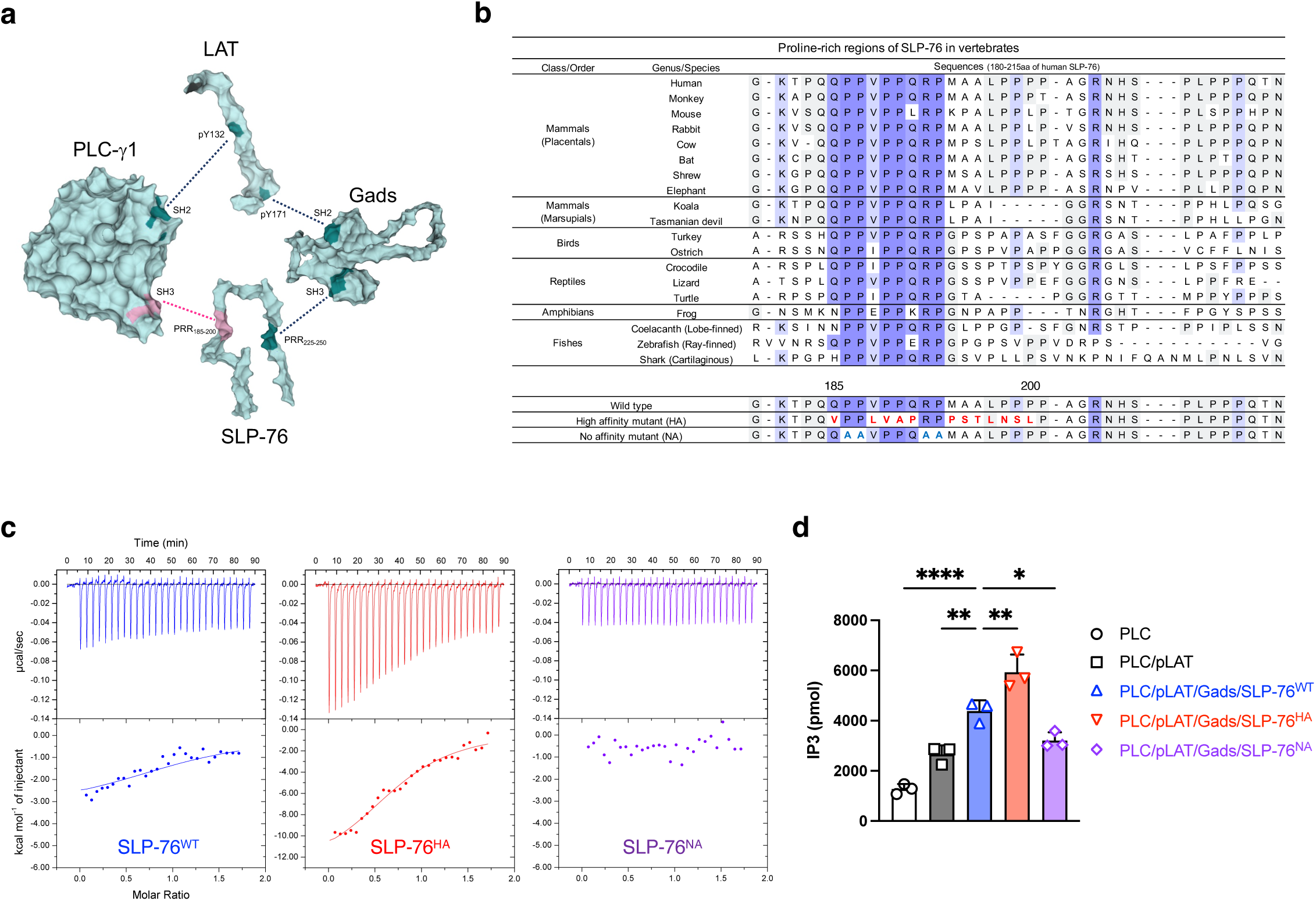
*In vitro* reconstitutional characterization of the tetrameric protein complex containing LAT, Gads, SLP-76 and PLC-γ1. **a**, Schematic of tetrameric structure (dark cyan, strong interaction sites; dark red, weak interaction site between PLC-γ1 and SLP-76). The 3D structure of PLC-γ1 is based on PDB:6PBC^47, 48^. The 3D model structure of LAT (aa 90-193), Gads (aa 1-330), and SLP-76 (aa 103-258) was generated by Alphafold3. **b**, Sequence alignment of the proline-rich region in SLP-76 among 19 jawed vertebrates. Multiple residues shown in red are substituted amino acids in the high affinity mutant. Four alanines in cyan are substituted amino acids in the no affinity mutant. **c**, Spectrum of isothermal titration calorimetry for the interaction between PLC-γ1 and SLP-76^WT^ (wild-type), SLP-76^HA^ (high affinity mutant), and SLP-76^NA^ (no affinity mutant), measured by VP-ITC. **d**, The levels of IP3 determined by PLC-γ1 catalytic activity assay. Data were generated from 3 independent experiments. Statistics was determined by ordinary one-way ANOVA. *; p<0.05, **; p<0.01, ****; p<0.0001.

**Table 1:** Thermodynamic parameters of individual protein-protein interactions determining tetramer formation among LAT, Gads, SLP-76 and PLC-γ1 This thermodynamic parameter table combines data from our previous studies^3, 5^ and the current study. In all ITC analyses, LAT or SLP-76 was in the syringe and injected into PLC-γ1 or Gads in the cell. The standard deviations shown in the data from the current study were calculated from 3 independent experiments.

| Interactions (Protein 1 / Protein 2) | Size of protein 1 | Size of protein 2 | K<sub>d</sub> (nM) | Temp. (°C) | ΔH (kcal/mol) | -TΔS (kcal/mol) | ΔG (kcal/mol) | Reference |
| --- | --- | --- | --- | --- | --- | --- | --- | --- |
| PLC-γ1 / LAT | 550-846 | 122-140 <sup>pY132</sup> | 62 | 25 | -15.7 ± 0.3 | 6.0 ± 0.3 | -9.8 ± 0.2 | * |
|  | 1-1219 | 125-178 <sup>pY132, pY171</sup> | 84 | 20 | -5.4 | -4.1 | -9.5 | ** |
| Gads / LAT | 1-322 | 161-180 <sup>pY171</sup> | 164 | 25 | -9.6 ± 0.6 | 0.3 ± 0.5 | -9.2 ± 0.1 | * |
|  | 1-322 | 125-178 <sup>pY132, pY171</sup> | 213 | 20 | -10.7 | 1.7 | -9.0 | ** |
| Gads / SLP-76 | 1-322 | 157-420 (WT) <sup>1</sup> | 19 | 25 | -12.0 ± 1.7 | 1.7 ± 1.8 | -10.4 ± 0.3 | * |
|  | 1-322 | 159-533 (WT) | 3.6 | 20 | -15.0 | 4.6 | -10.4 | ** |
| PLC-γ1 / SLP-76 | 550-846 | 157-420 (WT) | ND | 25 | ND | ND | ND | * |
|  | 550-846 | 157-420 (WT) | 3345 | 4 | -1.8 ± 0.1 | -5.6 ± 0.1 | -7.3 ± 0.0 | * |
|  | 1-1219 | 159-533 (WT) | ND | 20 | ND | ND | ND | ** |
| PLC-γ1 / SLP-76 | 1-1219 | 103-258 (WT) | 2317 ± 853 | 5 | -3.8 ± 1.0 | -3.4 ± 1.2 | -7.2 ± 0.3 | *** |
|  | 1-1219 | 103-258 (HA) <sup>2</sup> | 895 ± 100 | 5 | -10.9 ± 2.3 | 3.2 ± 2.3 | -7.7 ± 0.1 | *** |
|  | 1-1219 | 103-258 (NA) <sup>3</sup> | ND | 5 | ND | ND | ND | *** |
|  | 1-1219 | 185-200 (WT) | 3989 ± 1712 | 5 | -3.3 ± 0.7 | -3.6 ± 0.9 | -6.9 ± 0.3 | *** |
|  | 1-1219 | 185-200 (HA) | 1116 ± 389 | 5 | -10.3 ± 1.2 | 2.7 ± 1.3 | -7.6 ± 0.3 | *** |
|  | 1-1219 | 185-200 (NA) | ND | 5 | ND | ND | ND | *** |

1. Wild Type (WT) 2. High Affinity mutant (HA) 3. No Affinity mutant (NA)
\* Houtman et al., *Biochemistry* (2004) \*\* Manna et al., *PNAS* (2018) \*\*\* This study

To evaluate how much an alternation of the affinity between SLP-76 and PLC-γ1 could modulate the activation of PLC-γ1 in the context of the tetrameric complex, we designed two SLP-76 mutants and compared them to wild-type SLP-76 (SLP-76^WT^) in the formation of the tetrameric complex and PLC-γ1 activation. The sequence changes employed to create the high affinity (SLP-76^HA^) and no affinity mutant of SLP-76 (SLP-76^NA^) were indicated in Fig. 1b. The sequence of SLP-76^HA^ was obtained from previous screening data for the ligand preferences of the SH3 domain of PLC-γ1 using a phage-display library^7^. SLP-76^NA^ was newly designed as a negative mutant based on the alignment and previous research^8^. Purity of the three SLP-76 polypeptides was confirmed by SDS-PAGE (Extended Data Fig. 1b). We performed the ITC assay at 5°C to compare the thermodynamic parameters of binding to full length PLC-γ1 between the WT and SLP-76 PRR mutants (Fig. 1c and Table 1). The interaction of the SLP-76^WT^ polypeptide (aa 103-258) with PLC-γ1 had a K_d_ value of 2317 nM, and produced a small heat change (–3.8 kcal/mol), which was consistent with our previous study^3, 5^. K_d_ value differences between our previous and current studies are likely because we now used full length PLC-γ1 instead of the PLC-γ1 SH3 domain alone. The affinity of SLP-76^HA^ was 895 nM with a heat change (–10.9 kcal/mol), indicating that the high affinity mutation strengthens the binding of SLP-76 to PLC-γ1 and suggesting that this decrease in the K_d_ value is due to an increased change in enthalpy (ΔH). The SLP-76^NA^ showed no interaction with PLC-γ1, and consequently the K_d_ value could not be determined. We also employed a set of short peptides from the SLP-76 PRR (aa 185-200). Their interaction with PLC-γ1 was of similar but lower affinity than the longer SLP-76_103-258_ polypeptides (Table 1, Extended Data Fig. 1c).

It was previously reported that the SH3 domains of Itk and Lck also interact with the SLP-76 PRR^8, 9, 10, 11, 12, 13^. We performed sequence alignments of regions of the three SH3 domains, Itk_171-231_, Lck_61-121_ and, as above, PLC-γ1_791-851_ and found that the former two sequences lacked two of the four amino acids (indicated in red) critical for binding to the SLP-76 PRR (Extended Data Fig. 1d). We then carried out ITC assays to assess if this region of the Itk SH3 domain might bind to the three different forms of the SLP-76 PRR_185-200_ peptides and found that full-length Itk was unable to interact with the SLP-76 PRR (Extended Data Fig. 1e,f). The conserved proline-rich PIPRSP motif in LAT_54-90_ has recently been shown to associate with the Lck SH3 domain^14^. Our sequence alignment analysis revealed no similar feature between that LAT proline-rich motif and SLP-76 PRR_180-207_ (Extended Data Fig. 1g). Under the conditions of our analysis, we concluded that the SH3 domain of PLC-γ1 would be the major binding partner of the SLP-76 PRR_185-200_.

Next, we assessed the impact of the three SLP-76 PRR sequences on activation of PLC-γ1 *in vitro* using liposomes containing PIP2, the PLC-γ1 substrate^6^ (Fig. 1d). Recruitment of PLC-γ1 to liposomes containing phospho-LAT alone, proteins interacting with a K_d_ of 62 nM, caused an activation of PIP2 hydrolysis of 2.1-fold compared with liposomes incubated with PLC-γ1 alone (Fig. 1d: white vs. gray). This enhancement was consistent with our previous reconstitution study^6^. Reconstitution of the LAT-Gads-SLP-76^WT^-PLC-γ1 tetramer further enhanced PLC-γ1 activity on PIP2-bearing liposomes by 1.7-fold, indicating the importance of the SLP-76-PLC-γ1 interaction in the tetramer despite its low affinity with a K_d_ of 2317 nM (Fig. 1d: gray vs. blue). Substitution of SLP-76^WT^ with SLP-76^HA^ further increased the activity by 1.4-fold, and the tetramer with SLP-76^HA^ activated the function of PLC-γ1 by 4.6-fold in total (Fig. 1d: blue vs. red and white vs. red, respectively). In contrast, substitution with SLP-76^NA^ decreased the activity to that induced in the presence of PLC-γ1 and phospho-LAT without the Gads-SLP-76 dimer (Fig. 1d: gray vs. purple). These results confirm that the interaction of the SLP-76 PRR and the PLC-γ1 SH3 domain regulate PLC-γ1. Enhancing the affinity of the SLP-76-PLC-γ1 interaction likely leads to an increase in the frequency and/or duration of the closed circular tetramer formation, thereby enhancing PLC-γ1 activity.

### Generation of mice with conditional introduction of the SLP-76^HA^ mutation in a T cell lineage-specific manner

To evaluate the biological significance of the weak interaction between SLP-76 and PLC-γ1 in T cells under physiological settings, we created a mouse line in which the WT SLP-76 PRR_185-200_ was conditionally replaced with the SLP-76^HA^ (Fig. 2a and Extended Data Fig. 2a). A targeting vector was constructed consisting of WT SLP-76 cDNA spanning exons 8 through 21 (SLP-76 cDNA Ex. 8-21) followed by a polyadenylation (poly A) sequence, which was flanked by LoxP sequences. The second LoxP sequence was followed by the mutated exon 8 encoding the SLP-76^HA^. Using CRISPR/Cas9-mediated gene editing, the WT exon 8 of the *Lcp2* gene (encoding the SLP-76 protein) was replaced with the targeting vector. One founder mouse (C1) carrying the successfully recombineered *Lcp2^L^*^8^*^:21L-mE^*^8^ allele was identified by PCR using GT-1 and GT-2 primer sets (Extended Data Fig. 2b,c). As expected, the WT mouse showed neither GT-1 nor GT-2 PCR product (Extended Data Fig. 2c). The transcription of the *Lcp2^L^*^8^*^:21L-mE^*^8^ allele terminated at the poly A sequence without Cre recombinase activity, thus producing WT SLP-76 protein. Lineage-specific promoter/enhancer-driven Cre excised the LoxP-flanked SLP-76 cDNA Ex. 8-21 and poly A sequence, causing the mutated exon 8 and the exons 9 through 21 to follow the exon 7 (*Lcp2^mE^*^8^ allele). This recombineered allele produced full-length SLP-76 protein carrying the SLP-76^HA^ (Fig. 2a). Because lineage-specific Cre-mediated gene manipulation is not always precise and/or complete^15^, Lcp2^L8:21L-mE8^ mice were mated with *Rosa26^LSL-eYFP^* fate-mapping reporter (R26R-EYFP) mice to induce YFP expression in cells with Cre activity, thus marking the mutated SLP-76-expressing cells^16^ (Fig. 2a).

**Figure 2:**
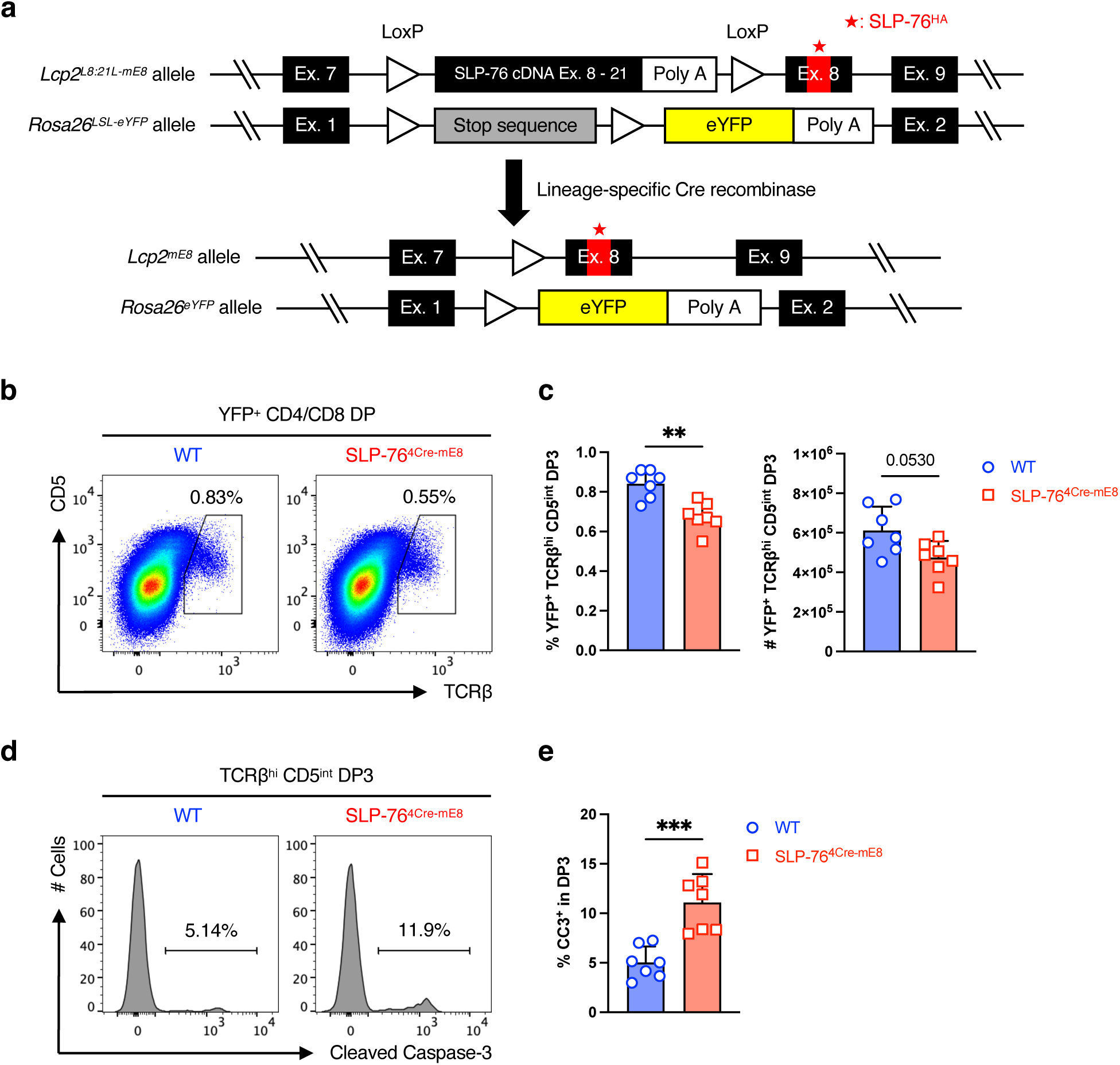
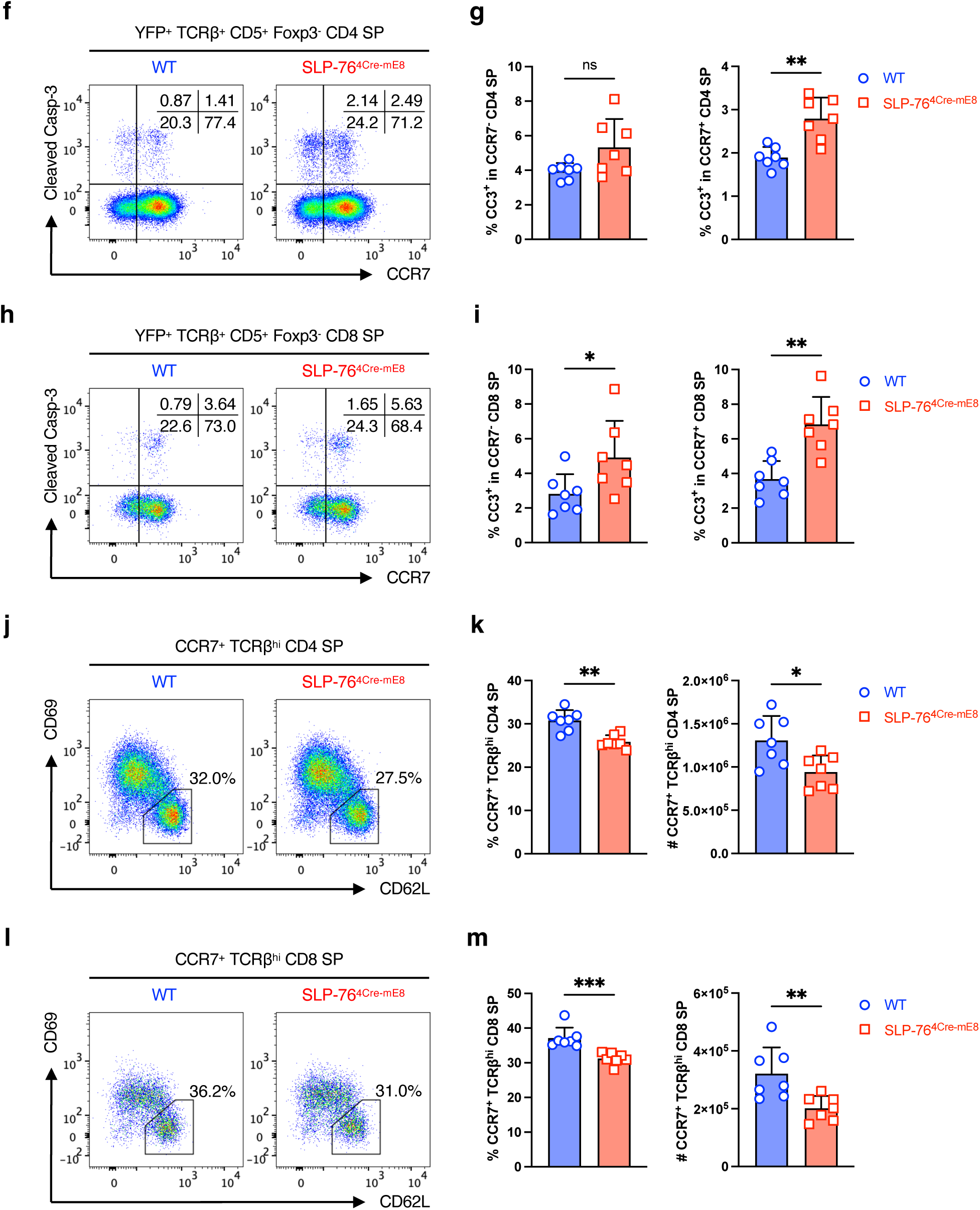
Altered thymocyte development caused by the SLP-76^HA^ mutation. **a**, Diagram of lineage-specific Cre-mediated expression of SLP-76 carrying the SLP-76^HA^ mutation and YFP fate-mapping reporter in mice. **b**, Representative pseudocolor plot analysis of YFP^+^ CD4/CD8 DP thymocytes for a TCRβ^high^CD5^int^ DP3 subset. **c**, Statistical analysis for the frequency (left) and absolute number (right) of DP3 subset (n=7). **d**, Representative histogram analysis in DP3 thymocytes for the frequency of cleaved Caspase-3 (CC3) expression. **e**, Statistical analysis for the frequency of CC3^+^ DP3 thymocytes (n=7). **f**, Representative pseudocolor plot analysis of YFP^+^TCRβ^+^CD5^+^ CD4 SP thymocytes for CC3 expression. The numbers indicate the frequency of cells in each quadrant. **g**, Statistical analysis for the frequency of CC3^+^ cells in CCR7^-^ (left) and CCR7^+^ (right) CD4 SP thymocytes (n=7). **h**, Representative pseudocolor plot analysis of YFP^+^TCRβ^+^CD5^+^ CD8 SP thymocytes for CC3 expression. The numbers indicate the frequency of cells in each quadrant. **i**, Statistical analysis for the frequency of CC3^+^ cells in CCR7^-^ (left) and CCR7^+^ (right) CD8 SP thymocytes (n=7). **j**, Representative pseudocolor plot analysis of CCR7^+^TCRβ^high^ CD4 SP thymocytes for CD62L and CD69 expressions. **k**, Statistical analysis for the frequency (left) and absolute number (right) of CD62L^+^CD69^-^ most mature (M2) medullary CD4 SP thymocytes (n=7). **l**, Representative pseudocolor plot analysis of CCR7^+^TCRβ^high^ CD8 SP thymocytes for CD62L and CD69 expressions. **m**, Statistical analysis for the frequency (left) and absolute number (right) of CD62L^+^CD69^-^ most mature (M2) medullary CD8 SP thymocytes (n=7). Statistics was determined by two-sided Student’s t-test. ns; not significant, *; p<0.05, **; p<0.01, ***; p<0.001.

### Altered thymocyte development caused by the SLP-76^HA^ mutation

To examine the effect of the SLP-76^HA^ mutation on T cell development in the thymus, R26R-EYFP mice and Lcp2^L8:21L-mE8^ x R26R-EYFP mice were crossed onto CD4^Cre^ mice^17^. At 8-12 weeks of age, total thymocyte profiles including the frequency and the absolute number of double-negative (DN), CD4/CD8 double-positive (DP), CD4 single-positive (SP), and CD8 SP cells were grossly normal in Lcp2^L8:21L-mE8^ x R26R-EYFP x CD4^Cre^ (SLP-76^4Cre-mE8^) mice, with a slight increase in the frequency of DN thymocytes (Extended Data Fig. 2d,e). However, SLP-76^4Cre-mE8^ YFP^+^ DP thymocytes showed a slight but significant decrease in the frequency of post-positive selection TCRβ^high^CD5^int^ DP3 cells^18^ (Fig. 2b,c). This finding was accompanied by an increased frequency of DP3 thymocytes expressing cleaved Caspase-3 (CC3) in SLP-76^4Cre-mE8^ mice (Fig. 2d,e). These results indicate that the SLP-76^HA^ mutation promotes the apoptosis of DP thymocytes that normally survive after positive selection, presumably due to enhanced TCR signal strength during the recognition of MHC.

Next, we assessed whether the SLP-76^HA^ mutation would affect negative selection of CD4 and CD8 SP thymocytes. YFP^+^TCRβ^+^CD5^+^Foxp3^-^ conventional CD4 and CD8 SP thymocytes that had experienced TCR recognition of self-peptide/MHC were subdivided on the basis of CCR7 expression. CCR7 is required for positively selected CD4 and CD8 SP thymocytes to migrate from cortex to medulla, a major niche for negative selection^19, 20^. SLP-76^4Cre-mE8^ mice showed a slight but not significant increase in the frequency of CC3^+^ cells in CCR7^-^ CD4 SP thymocytes but had a significant enhancement of that in CCR7^+^ CD4 SP thymocytes (Fig. 2f,g and Extended Data Fig. 2f). Moreover, the frequency of CC3^+^ cells was increased in both CCR7^-^ and CCR7^+^ CD8 SP thymocytes from SLP-76^4Cre-mE8^ mice (Fig. 2h,i and Extended Data Fig. 2g). Consistent with these results, SLP-76^4Cre-mE8^ mice showed a significant decrease in the frequency and absolute number of CCR7^+^TCRβ^high^CD62L^+^CD69^-^ medullary CD4 and CD8 SP most mature (M2) thymocytes that had completed negative selection^21, 22^ (Fig. 2j–m). These results imply that the SLP-76^HA^ mutation promotes the apoptosis of CD4 and CD8 SP thymocytes that normally pass the negative selection step, presumably due to enhanced TCR signal strength during the recognition of self-peptide/MHC.

### Enhancement of memory-phenotype T cells in the periphery of SLP-76^4Cre-mE^^8^ mice

We assessed whether the SLP-76^HA^ mutation would alter the phenotype of peripheral CD4^+^ and CD8^+^ T cells in pooled whole lymph nodes (LNs) (axillary, brachial, superficial/deep cervical, inguinal, and mesenteric) of 8–12-week-old mice in a steady state. CD3^+^YFP^+^CD4^+^ T cells were subdivided into 3 populations based on Foxp3 and Helios expressions: Foxp3^-^ conventional T cells, Foxp3^+^Helios^+^ thymic-derived regulatory T cells (tTregs), and Foxp3^+^Helios^-^ peripherally induced regulatory T cells (pTregs)^23^. The frequency and number of all 3 populations were unaffected in SLP-76^4Cre-mE8^ mice (Fig. 3a,b and Extended Data Fig. 3a,b). Moreover, CD44^low^CD62L^high^ central Tregs (cTregs) and CD44^high^CD62L^low^ effector Tregs (eTregs) in both tTregs and pTregs were comparable between WT and SLP-76^4Cre-mE8^ mice, except a slight increase in the number of eTregs in tTregs^24, 25, 26^ (Fig. 3c–f and Extended Data Fig. 3c,d). These results imply that the SLP-76^HA^ mutation barely influences the homeostasis and functional property of Tregs. Subsequent analysis of conventional CD4^+^ T cells revealed that SLP-76^4Cre-mE8^ mice had a mild increase in the frequency of CD44^high^CD62L^low^ memory-phenotype T (T_MP_) cells that arise from naïve CD4^+^ T cells upon recognition of a self-peptide/MHC II complex^27^ (Fig. 3g,h and Extended Data Fig. 3a). Additionally, SLP-76^4Cre-mE8^ mice had a marked increase in the frequency and absolute number of CD44^high^Ly-6C^high^ CD8^+^ T_MP_ cells that also expressed CD62L and CD122/IL-2Rβ chain^28, 29^ (Fig. 3i,j and Extended Data Fig. 3e–h). These results imply that when naïve CD4^+^ and CD8^+^ T cells recognize self-peptide/MHC complexes in the secondary lymphoid organs, enhanced TCR signal strength as a consequence of the SLP-76^HA^ mutation leads to increased generation of T_MP_ cells.

**Figure 3:**
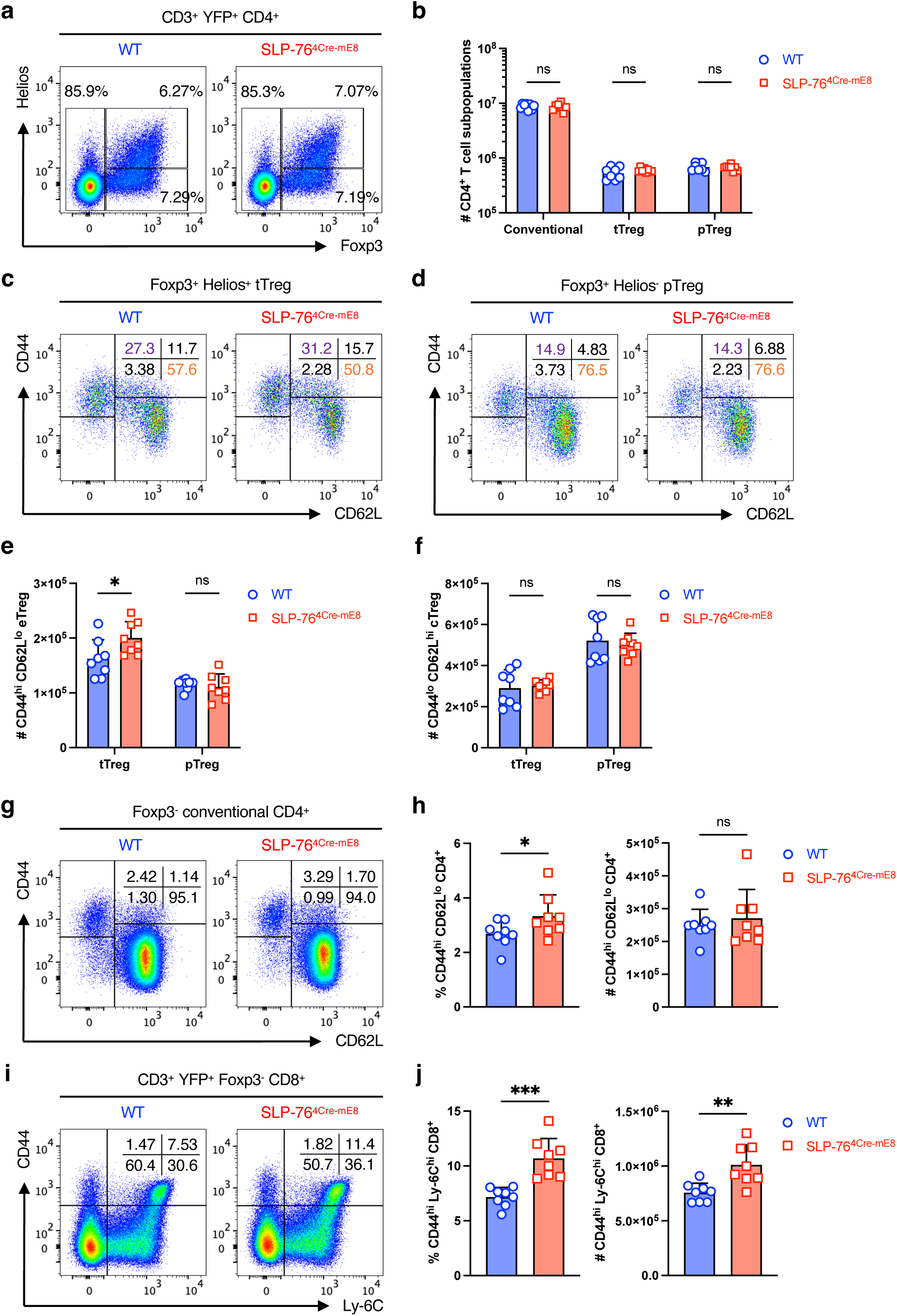
Characterization of peripheral T cells in SLP-76^4Cre-mE8^ mice. **a**, Representative pseudocolor plot analysis of CD3^+^YFP^+^CD4^+^ T cells from pooled whole LNs for Foxp3 and Helios expressions. **b**, Statistical analysis for the absolute number of Foxp3^-^ conventional CD4^+^ T cells, Foxp3^+^Helios^+^ thymic-derived Tregs (tTregs), and Foxp3^+^Helios^-^ peripherally induced Tregs (pTregs) (n=8). **c**,**d**, Representative pseudocolor plot analysis of tTregs (**c**) and pTregs (**d**) for CD44 and CD62L expressions. The numbers indicate the frequency of cells in each quadrant. **e**,**f**, Statistical analysis for the absolute number of CD44^high^CD62L^low^ effector Tregs (eTregs) (**e**) and CD44^low^CD62L^high^ central Tregs (cTregs) (**f**) in tTreg and pTreg compartments. **g**, Representative pseudocolor plot analysis of conventional CD4^+^ T cells for CD62L and CD44 expressions. **h**, Statistical analysis for the frequency (left) and absolute number (right) of CD44^high^CD62L^low^ memory-phenotype CD4^+^ T (T_MP_) cells (n=8). **i**, Representative pseudocolor plot analysis of conventional CD8^+^ T cells for Ly-6C and CD44 expressions. The numbers indicate the frequency of cells in each quadrant. **j**, Statistical analysis for the frequency (left) and absolute number (right) of CD44^high^Ly-6C^high^ CD8^+^ T_MP_ cells (n=8). Statistics was determined by two-way ANOVA (**b**,**e**,**f**) and two-sided Student’s t-test (**h**,**j**). ns; not significant, *; p<0.05, **; p<0.01, ***; p<0.001.

### SLP-76^HA^ mutation enhances the responsiveness of OT-I CD8^+^ T cells to OVA altered peptide ligands with lower affinity in vitro and in vivo

We next examined whether the SLP-76^HA^ mutation would influence the degree of activation of peripheral T cells. To this end, Lcp2^L8:21L-mE8^ x R26R-EYFP mice were crossed onto hCD2^Cre^ mice, which excises the LoxP-flanked SLP-76 cDNA Ex. 8-21 and the poly A sequence after negative selection, thus circumventing the effect of the SLP-76^HA^ mutation on thymocyte development^30^. The resultant mice were further crossed onto OT-I TCR transgenic mice, to enable use of the variety of OVA_257-264_ altered peptide ligands (APLs) with various affinity for an OT-I TCR to evaluate the strength of TCR-mediated signaling in CD8^+^ T cells^31^ ^32^. R26R-EYFP x hCD2^Cre^ x OT-I (WT OT-I) mice and Lcp2^L8:21L-mE8^ x R26R-EYFP x hCD2^Cre^ x OT-I (SLP-76^2Cre-mE8^ OT-I) mice were bred on a CD45.1 and CD45.2 background, respectively, to distinguish OT-I CD8^+^ T cells of the two different genotypes by flow cytometry^33^.

First, we assessed the effect of the SLP-76^HA^ mutation on early activation of OT-I CD8^+^ T cells in response to OVA-derived peptides *in vitro*. CD8^+^ T cells isolated from WT OT-I CD45.1 and SLP-76^2Cre-mE8^ OT-I CD45.2 mice were mixed at a 1:1 ratio and co-cultured with T cell-depleted spleens (TdS) cells from (CD45.1 x CD45.2) F1 mice that had been loaded with WT OVA peptide N4 (SII<u>N</u>FEKL; ligand potency of 1) or 2 different OVA APLs with lower affinities for OT-I TCR, Q4H7 (SII<u>Q</u>FE<u>H</u>L; ligand potency of 1/167) and G4 (SII<u>G</u>FEKL; ligand potency of 1/7515)^32^. Twenty hours later, activated OT-I CD8^+^ T cells were analyzed for expression of CD69 and PD-1 by flow cytometry (Extended Data Fig. 4a). When stimulated with N4 OVA peptide, SLP-76^2Cre-mE8^ OT-I cells expressed the same levels of CD69 (combined frequency of CD69^+^PD-1^-^ and CD69^+^PD-1^+^) as did WT OT-I cells at all concentrations of peptide (Fig. 4a and Extended Data Fig. 4b). The dose of N4 peptide inducing a 50% maximum of CD69 expression (EC_50_) was comparable between WT and SLP-76^2Cre-mE^^8^ OT-I cells in 3 independent experiments (Fig. 4d,e). In contrast, stimulation with Q4H7 led to a higher frequency of CD69 expression on SLP-76^2Cre-mE8^ OT-I cells at lower peptide concentrations ranging from 3.91 x 10^-10^ to 1.56 x 10^-9^ M than that on WT OT-I cells (Fig. 4b and Extended Data Fig. 4c). Q4H7-stimulated SLP-76^2Cre-mE^^8^ OT-I cells showed a 2-fold decrease in the EC_50_ of CD69 (Fig. 4d,e). Moreover, SLP-76^2Cre-mE8^ OT-I cells expressed more CD69 at any given concentrations of G4 (10^-8^ – 10^-5^ M), but the EC_50_ of CD69 could not be determined because of poor activation of both WT and SLP-76^2Cre-mE8^ OT-I cells (Fig. 4c and Extended Data Fig. 4d). These results indicate that the SLP-76^HA^ mutation does not affect the responsiveness of CD8^+^ T cells to a strong agonist peptide sufficient to induce robust T cell activation but renders them more responsive to weak agonists.

**Figure 4:**
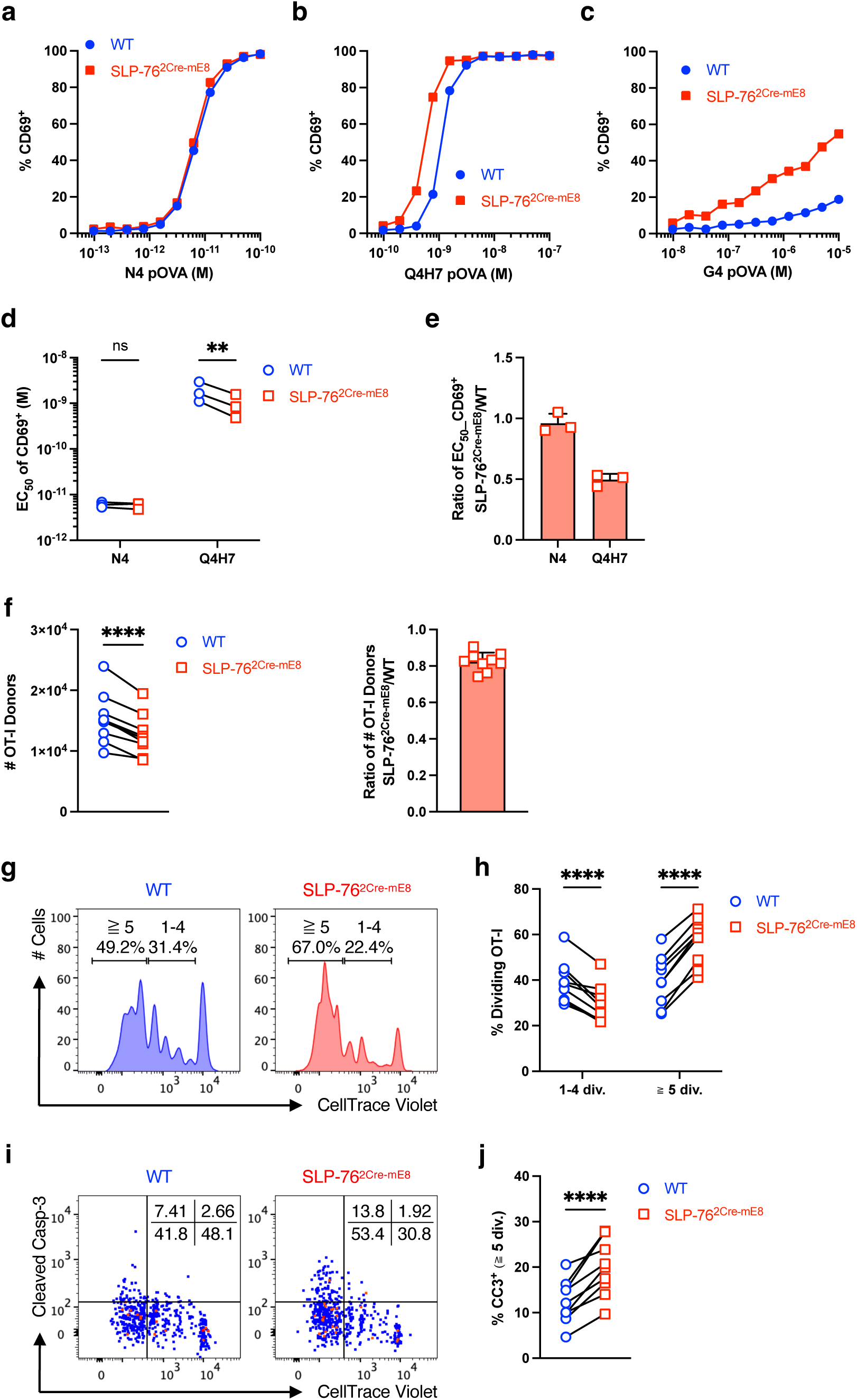
SLP-76^HA^ mutation enhances the responsiveness of OT-I CD8^+^ T cells to OVA altered peptide ligands (APLs) with lower affinity *in vitro* and *in vivo*. **a**–**e**, WT CD45.1 and SLP-76^2Cre-mE8^ CD45.2 OT-I CD8^+^ T cells were co-cultured with (CD45.1 x CD45.2) F1 TdS cells loaded with WT N4 OVA peptide or two different OVA APL (Q4H7 or G4) *in vitro* for 20 hours. Shown is the frequency of CD69^+^ WT (blue circle) and SLP-76^2Cre-mE8^ (red square) OT-I cells stimulated with varying concentrations of N4 (**a**), Q4H7 (**b**), and G4 (**c**) peptide. **d**,**e**, Statistical analysis for EC_50_ of CD69^+^ (**d**) and fold change in EC_50_ of CD69^+^ between WT and SLP-76^2Cre-mE8^ OT-I cells (**e**) in response to N4 or Q4H7 peptide from 3 independent experiments. **f**–**j**, CTV-labeled WT CD45.1 and SLP-76^2Cre-mE8^ CD45.2 OT-I CD8^+^ T cells were co-transferred into (CD45.1 x CD45.2) F1 mice. One day later, recipient mice were subcutaneously immunized with Q4H7/CFA. Donor OT-I cells in skin dLNs were analyzed at 3 days post-immunization by flow cytometry. **f**, Statistical analysis for the absolute number of WT and SLP-76^2Cre-mE8^ OT-I donor cells (left) and the ratio of SLP-76^2Cre-mE8^ to WT OT-I donor cells (right) (n=9). **g**, Representative histogram analysis for division of WT and SLP-76^2Cre-mE8^ OT-I donor cells. **h**, Statistical analysis for the frequency of OT-I donor cells that have undergone 1-4 divisions and those with more than 5 divisions (n=9). **i**, Representative pseudocolor plot analysis of WT and SLP-76^2Cre-mE8^ OT-I donor cells for cleaved Caspase-3 (CC3) expression. The numbers indicate the frequency of cells in each quadrant. **j**, Statistical analysis for CC3^+^ OT-I donor cells that have undergone more than 5 divisions (n=9). Statistics was determined by two-way ANOVA (**d**,**h**) and paired t-test (**f**,**j**). ns; not significant, **; p<0.01, ****; p<0.0001.

To examine the effect of the SLP-76^HA^ mutation on CD8^+^ T cell priming *in vivo*, CD8^+^ T cells isolated from WT OT-I CD45.1 and SLP-76^2Cre-mE8^ OT-I CD45.2 mice were mixed at a 1:1 ratio, labeled with CellTrace Violet (CTV), and transferred into (CD45.1 x CD45.2) F1 mice. One day later, recipient mice were subcutaneously immunized with Q4H7 APL emulsified in complete Freund’s adjuvant (CFA). We chose this APL because it led to enhanced responsiveness of SLP-76^2Cre-mE8^ OT-I cells *in vitro* compared with WT OT-I cells (Fig. 4b,d,e). Skin draining LNs (dLNs) (axillary, brachial, and inguinal) were excised at 3 days post-immunization to analyze donor OT-I cells by flow cytometry (Extended Data Fig. 4e). The absolute number of SLP-76^2Cre-mE8^ OT-I cells were slightly but significantly decreased, compared with WT OT-I cells from the same recipient mice (Fig. 4f). However, the proportion of SLP-76^2Cre-mE8^ OT-I cells that had undergone more than 5 divisions was substantially increased (Fig. 4g,h). Indeed, highly dividing SLP-76^2Cre-mE8^ OT-I cells had a significant increase in the frequency of CC3^+^ cells than did WT OT-I cells (Fig. 4i,j). These results indicate that the SLP-76^HA^ mutation accelerates the proliferation of CD8^+^ T cells primed with a weak agonist *in vivo* but renders highly dividing CD8^+^ T cells more apoptotic, thus decreasing the number of antigen-specific CD8^+^ T cells during the priming phase.

### SLP-76^HA^ mutation restrains the polarization of P14 CD8^+^ T cells toward T_CM_ cells in the memory phase upon acute LCMV infection

TCR signal strength has been demonstrated as one of the key determinants regulating a balance between effector memory (T_EM_) and central memory (T_CM_) CD8^+^ T cell generation in acute bacterial and viral infection models^34, 35, 36, 37, 38, 39^. To assess the T cell-autonomous effect of the SLP-76^HA^ mutation on the T_EM_/T_CM_ balance, CD8^+^ T cells isolated from WT P14 CD45.1 mice (R26R-EYFP x hCD2^Cre^ x P14 x CD45.1) and SLP-76^2Cre-mE8^ P14 CD45.2 mice (Lcp2^L8:21L-mE8^ x R26R-EYFP x hCD2^Cre^ x P14 x CD45.2) were mixed at a 1:1 ratio and transferred into (CD45.1 x CD45.2) F1 mice^40^. One day later, recipient mice were infected with the Armstrong strain of lymphocytic choriomeningitis virus (LCMV_Arm_)^41^. YFP^+^ P14 donor cells in the spleen were analyzed at 7 and 28 days post-infection by flow cytometry. The SLP-76^HA^ mutation slightly enhanced the number of CD127^-^KLRG1^+^ short-lived effector cells (SLEC) but had little effect on those of CD127^+^KLRG1^-^ memory-precursor effector cells (MPEC) at 7 days post-infection (Fig. 5a,b). At 28 days post-infection, the absolute number of SLP-76^2Cre-mE8^ P14 cells was equivalent to that of co-transferred WT P14 cells (Extended Data Fig. 5a,b). Notably, SLP-76^2Cre-mE8^ P14 cells showed a marked decrease in the frequency of CD127^+^KLRG1^-^ T_CM_ cells with a concomitant increase in that of CD127^-^KLRG1^+^ T_EM_ cells (Fig. 5c,d). Consistent with these results, the frequency of a TCF1^+^KLRG1^-^ population corresponding to T_CM_ cells was markedly decreased in SLP-76^2Cre-mE^^8^ P14 donor cells^42^ (Fig. 5e,f). The number of SLP-76^2Cre-mE^^8^ P14 T_EM_ cells was slightly increased by 1.3-fold, but this increase was not statistically significant. In contrast, there was a 2-fold decrease in the number of SLP-76^2Cre-mE^^8^ P14 T_CM_ cells (Fig. 5g,h). These results indicate that the SLP-76^HA^ mutation does not affect the skewing toward MPEC in the expansion phase but hinders the generation of T_CM_ cells in the early memory phase in a T cell-autonomous manner. The same trend was evident when splenic CD8^+^ T cells bound to LCMV_gp33-41_/H-2D^b^ tetramer (gp33-Tet) were analyzed for T_EM_ and T_CM_ cells in SLP-76^2Cre-mE^^8^ mice bearing endogenous TCR repertoire that had been infected with LCMV_Arm_ for 30 days (Extended Data Fig. 5e,f). However, the absolute numbers of gp33-Tet^+^ T_EM_ and T_CM_ cells were not significantly different between WT and SLP-76^2Cre-mE8^ mice because of a large variation in the robustness of gp33-Tet^+^ CD8^+^ T cell responsiveness in both groups (Extended Data Fig. 5c,d,g).

**Figure 5:**
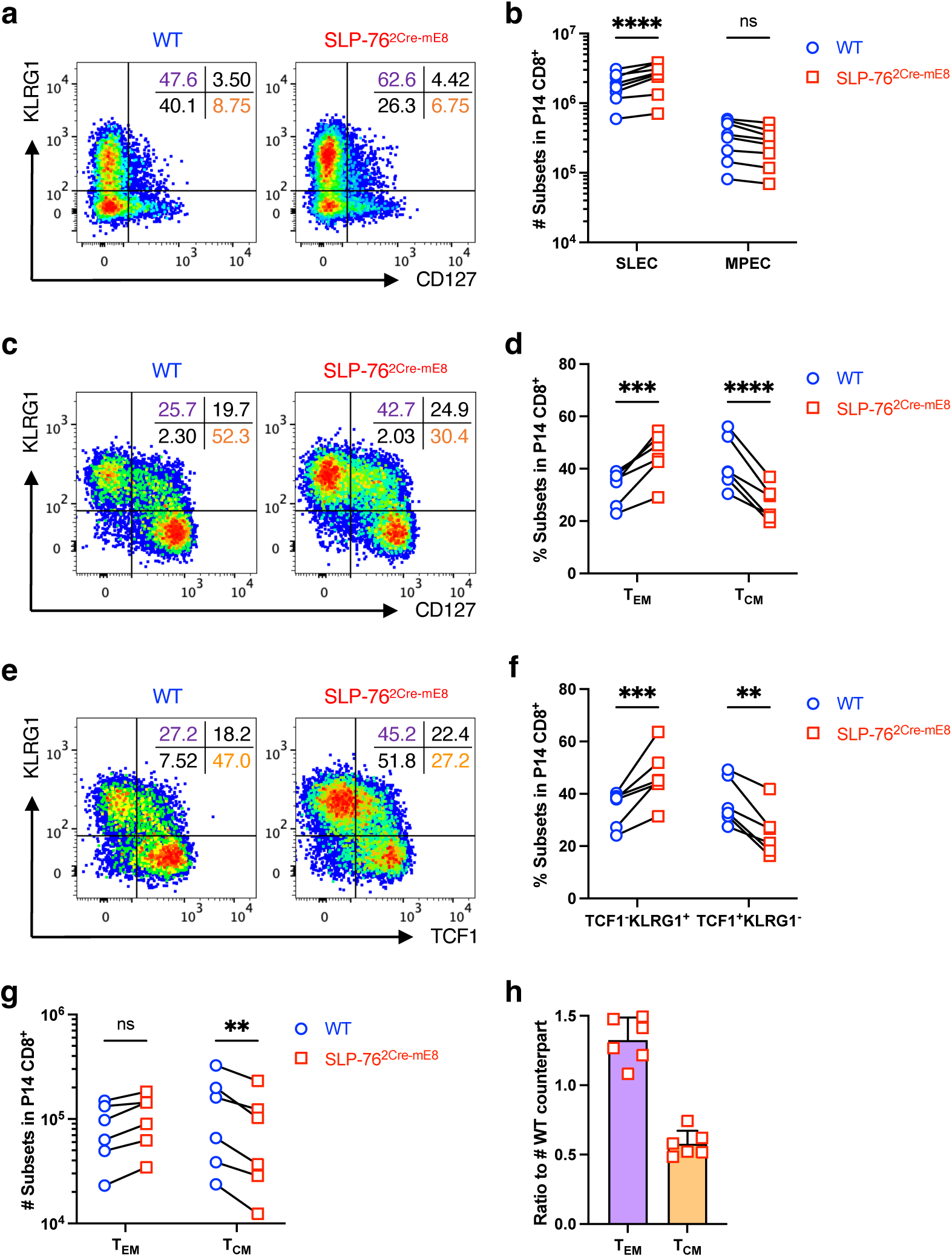
SLP-76^HA^ mutation restrains the polarization of P14 CD8^+^ T cells toward T_CM_ cells in the memory phase upon acute LCMV infection. WT CD45.1 and SLP-76^2Cre-mE8^ CD45.2 P14 CD8^+^ T cells were co-transferred into (CD45.1/CD45.2) F1 mice. One day later, the recipients were infected with LCMV_Arm_. **a**,**b**, Seven days post-infection. **a**, Representative pseudocolor plot analysis of WT and SLP-76^2Cre-mE8^ P14 donor cells for CD127 and KLRG1 expressions. The numbers indicate the frequency of cells in each quadrant. **b**, Statistical analysis for the absolute number of SLEC (CD127^-^KLRG1^+^) and MPEC (CD127^+^KLRG1^-^) in P14 donor cells (n=8). **c**–**h**, Twenty-eight days post-infection. **c**, Representative pseudocolor plot analysis of WT and SLP-76^2Cre-mE8^ P14 donor cells for CD127 and KLRG1 expressions. The numbers indicate the frequency of cells in each quadrant. **d**, Statistical analysis for the frequency of T_EM_ (CD127^-^KLRG1^+^) and T_CM_ (CD127^+^KLRG1^-^) P14 donor cells (n=6). **e**, Representative pseudocolor plot analysis of WT CD45.1 and SLP-76^2Cre-mE8^ P14 donor cells for TCF1 and KLRG1 expressions. The numbers indicate the frequency of cells in each quadrant. **f**, Statistical analysis for the frequency of TCF1^-^KLRG1^+^ and TCF1^+^KLRG1^-^ P14 donor cells (n=6). **g**, Statistical analysis for the absolute number of T_EM_ and T_CM_ P14 donor cells (n=6). **h**, Ratio of SLP-76^2Cre-mE^^8^ T_EM_ and T_CM_ cells to WT P14 counterparts. Statistics was determined by two-way ANOVA (**b**,**d**,**f**,**g**). ns; not significant, **; p<0.01, ***; p<0.001, ****; p<0.0001.

### SLP-76^HA^ mutation leads to the diminution in mature Tfh cells

T follicular helper (Tfh) cells are a subset of CD4^+^ helper T (Th) cells that supports humoral immune responses to T-dependent antigens by B cells in the germinal center (GC)^43^. As TCR signal strength has been proposed to play a crucial role in the fate decision of naïve CD4^+^ T cells toward either Tfh cells or non-Tfh cells^44, 45^, we assessed the T cell-autonomous effect of the SLP-76^HA^ mutation on Tfh differentiation. CD4^+^ T cells isolated from WT OT-II CD45.1 mice (R26R-EYFP x hCD2^Cre^ x OT-II x CD45.1) and SLP-76^2Cre-mE8^ OT-II CD45.2 mice (Lcp2^L8:21L-mE8^ x R26R-EYFP x hCD2^Cre^ x OT-II x CD45.2) were mixed at a 1:1 ratio and transferred into (CD45.1 x CD45.2) F1 mice^46^. One day later, recipient mice were immunized with OVA protein in CFA. YFP^+^ OT-II donor cells in skin dLNs were analyzed by flow cytometry at 10 days post-immunization (Extended Data Fig. 6a). SLP-76^2Cre-mE8^ OT-II cells showed a slight but significant decrease in absolute number by 1.6-fold, compared with WT OT-II cells from the same recipient mice (Extended Data Fig. 6b). Notably, the frequencies of PD-1^high^CXCR5^high^ and Bcl-6^high^CXCR5^high^ mature Tfh (GC-Tfh) cells were selectively decreased, whereas those of PD-1^-^ CXCR5^-^ non-Tfh cells and PD-1^int^CXCR5^int^ pre-Tfh cells was either slightly increased or unchanged (Fig. 6a–d). The numbers of SLP-76^2Cre-mE8^ non-Tfh and pre-Tfh cells were slightly decreased to a degree equivalent to that of total SLP-76^2Cre-mE8^ OT-II donor cells, whereas SLP-76^2Cre-mE8^ mature Tfh cells were markedly decreased by 2.5-fold (Fig. 6e,f). Consistent with these results, the frequency of dead cells in a CXCR5^+^ fraction was substantially enhanced in SLP-76^2Cre-mE8^ OT-II donor cells at 6 and 8 days post-immunization (Fig. 6g,h and Extended Data Fig. 6c). These results indicate that the SLP-76^HA^ mutation does not prevent antigen-primed CD4^+^ T cells from the fate decision toward Tfh cells but instead leads to a mild decrease in the number of clonally expanded CD4^+^ T cells. Importantly, the SLP-76^HA^ mutation results in the failure to maintain mature Tfh cells, presumably because the enhanced TCR signal strength caused by this mutation may promote death of pre-Tfh cells during cognate interaction with B cells.

**Figure 6:**
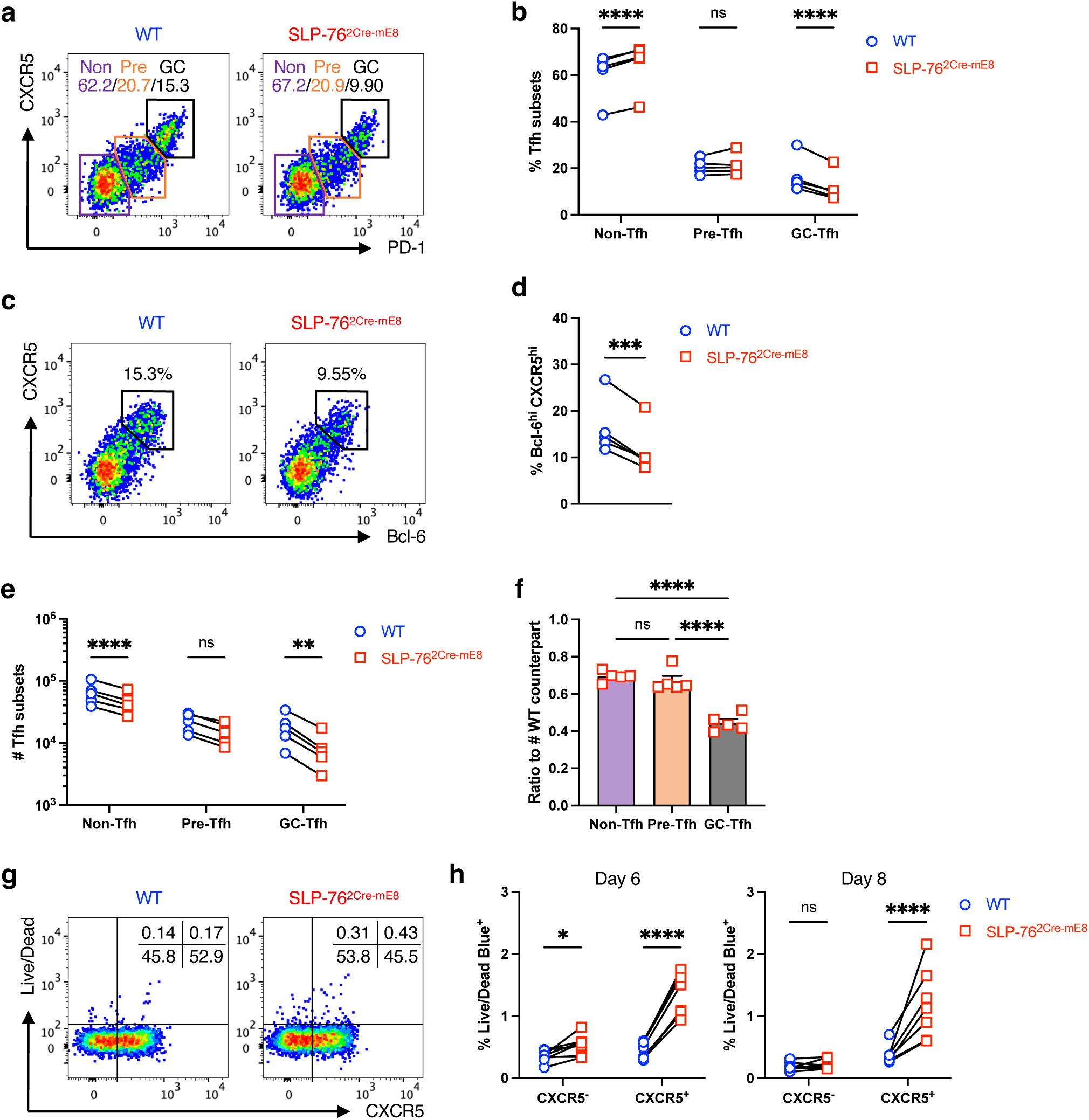
SLP-76^HA^ mutation leads to the diminution in mature Tfh cells. WT CD45.1 and SLP-76^2Cre-mE8^ CD45.2 OT-II CD4^+^ T cells were co-transferred into (CD45.1x CD45.2) F1 mice. One day later, recipient mice were subcutaneously immunized with EndoFit OVA in CFA. OT-II donor cells in skin dLNs were analyzed at 10 days post-immunization by flow cytometry. **a**, Representative pseudocolor plot analysis of WT and SLP-76^2Cre-mE8^ OT-II donor cells for PD-1 and CXCR5 expressions. The numbers indicate the frequency of cells in each population. **b**, Statistical analysis for the frequency of non-Tfh (PD-1^-^CXCR5^-^), pre-Tfh (PD-1^int^CXCR5^int^), and mature (GC) Tfh (PD-1^high^CXCR5^high^) OT-II donor cells. **c**, Representative pseudocolor plot analysis of WT and SLP-76^2Cre-mE8^ OT-II donor cells for Bcl-6 and CXCR5 expressions. **d**, Statistical analysis for the frequency of Bcl-6^high^CXCR5^high^ OT-II donor cells (n=5). **e**, Statistical analysis for the absolute number of non-Tfh, pre-Tfh, and GC-Tfh OT-II donor cells (n=5). **f**, Ratio of SLP-76^2Cre-mE8^ non-Tfh, pre-Tfh, and GC-Tfh cells to WT OT-II counterparts. **g**, Representative pseudocolor plot analysis of WT and SLP-76^2Cre-mE8^ OT-II donor cells for CXCR5 and Live/Dead Blue staining at 6 days post-immunization. The numbers indicate the frequency of cells in each population. **h**, Statistical analysis for the frequency of dead cells in CXCR5^-^ and CXCR5^+^ populations at 6 days (n=8) and 8 days (n=7) post-immunization. Statistics was determined by two-way ANOVA (**b**,**e**,**h**), paired t-test (**d**), and ordinary one-way ANOVA (**f**). ns; not significant, *; p<0.05, ***; p<0.001, ****; p<0.0001.

Taken together, the affinity of interaction between the SLP-76 PRR_185-200_ and the PLC-γ1 SH3 domain, which defines the level of PLC-γ1 activation induced upon TCR engagement, has a critical role in fine-tuning TCR signal strength in multiple settings. The native sequence and the low affinity of the interaction prevent the excessive death of thymocytes during positive and negative selection, restrains over-activation of CD8^+^ T cells leading to enhanced apoptosis during the priming period, limits the polarization toward CD8^+^ T_EM_ cells upon acute virus infection, and maintains normal levels of mature Tfh cells.

## Discussion

Activation of phospholipases are central to induction of many cellular changes in eukaryotic systems including proliferation, growth and migration. One of the multiple isoforms of phospholipase, PLC-γ1, is activated during the process of lymphocyte activation. This form of phospholipase C contains three regulatory domains, N-terminus SH2 (nSH2), C-terminus SH2 (cSH2), and SH3, that are important for its recruitment and activation^47, 48^. In T lymphocytes, antigen-induced activation through the TCR leads to phosphorylation of the LAT adapter molecule at its four distal tyrosine residues^1, 4^. The nSH2 domain of PLC-γ1 binds to phosphorylated Y132 of LAT. Many other adapter and signaling proteins bind to the three distal LAT phosphorylated tyrosine residues including a cytosolic dimer comprised of Gads and SLP-76. The proximity of PLC-γ1 and the Gads-SLP-76 dimer all bound to a LAT molecule enables formation of a tetrameric structure (LAT-Gads-SLP-76-PLC-γ1) in which the PLC-γ1 SH3 domain is able to bind to the SLP-76 PRR^2^.

Our laboratory subsequently identified the affinities of the domain interactions within this tetramer^5^. The LAT-Gads, Gads-SLP-76 and LAT-PLC-γ1 interactions are easily determined by ITC at room temperature. In contrast, the low affinity SLP-76-PLC-γ1 interaction can only be determined at 4°C. We then measured PLC-γ1 activation in an *in vitro* system using liposomes containing the PLC-γ1 substrate PIP2 and demonstrated that formation of the LAT-Gads-SLP-76-PLC-γ1 tetramer at the membrane results in the maximal induction of PLC-γ1 hydrolysis as measured by IP3 production^6^. This functional data in conjunction with our previous biophysical studies confirmed the importance of the SLP-76-PLC-γ1 interaction leading to circular tetramer formation. We proposed that formation of the circular tetramer would induce conformational changes in PLC-γ1 leading to its activation.

In the present study we first compared the amino acid sequences of SLP-76 and PLC-γ1 from multiple vertebrates to further test the significance of the low affinity SLP-76-PLC-γ1 interaction within the tetrameric complex. We confirmed that there was a high degree of sequence conservation in the two critical sites of interaction, the SLP-76 PRR_185-200_ and the PLC-γ1 SH3 domain, from human to fishes. These data suggest that the conserved low affinity interaction between these protein domains would play a critical role in modulating the circular tetrameric complex between open and closed conformations to optimize PLC-γ1 activation.

To test this hypothesis, we generated a set of recombinant SLP-76 proteins with the wild-type PRR_185-200_ sequence and with the SLP-76^NA^ and SLP-76^HA^ mutations predicted to lose and enhance the capacity to interact with PLC-γ1 via its SH3 domain, respectively. As predicted, our ITC analysis revealed that substitution of the native SLP-76 protein with the SLP-76^HA^ mutant resulted in enhanced binding to PLC-γ1, and that the SLP-76^NA^ mutant was unable to interact with PLC-γ1. We then performed the liposome-based assay to evaluate the effect of these sequence changes on PLC-γ1 activity. We found that incubation of the liposomes with the LAT-Gads-SLP-76^HA^-PLC-γ1 tetramer led to a small but significant increase in IP3 production compared to that with the LAT-Gads-SLP-76^WT^-PLC-γ1 tetramer. In contrast, formation of the tetramer containing the SLP-76^NA^ resulted in significantly lower levels of IP3 production than that with the SLP-76^WT^. These results indicate that the SLP-76-PLC-γ1 interaction is essential for regulation of the level of PLC-γ1 activation via formation of the closed circular tetramer.

The SLP-76 PRR_185-194_, identified as the minimal binding site for the PLC-γ1 SH3 domain, has also been reported to associate with the SH3 domain of Lck and Itk *in vitro* and *in vivo*^8, 9, 10, 11, 12,13^. These observations raise the possibility that the active SLP-76 PRR_185-200_ mutation used in our study might enhance the association of Lck and/or Itk within the TCR-induced multi-protein complex, thus increasing the activity of PLC-γ1 and the resultant TCR signal strength. However, this possibility can be excluded by the following pieces of evidence. First, sequence alignment of the Itk and Lck SH3 domains revealed that both of them lacked two of the four amino acids critical for binding to the SLP-76 PRR, which are present in the PLC-γ1 SH3 domain^8^. Second, our ITC analysis revealed no measurable interaction of Itk with any of the three forms of SLP-76 PRR_185-200_ peptides. Third, neither Lck nor Itk co-immunoprecipitates with SLP-76 in carefully controlled experiments using Lck-deficient J.CaM1 cells as a negative control, as demonstrated by Yablonski et al.^2^ Fourth, the SLP-76 PRR_180-207_ did not have any sequence feature similar to the PIPRSP motif in LAT_54-90_ that associates with the Lck SH3 domain^14^.

Although interaction of the Itk SH3 domain with the SLP-76 PRR is unlikely to occur, Itk binds to phosphorylated SLP-76 at Y145 via its SH2 domain with the adapter Nck and the enzyme Vav, thereby forming a heptameric signaling complex^49, 50, 51^. Activated Itk in turn phosphorylates SLP-76 at Y173, which primes PLC-γ1 for Itk-mediated phosphorylation of a regulatory tyrosine residue Y783^52, 53^. Structural studies have demonstrated how this phosphorylation event would result in conformation changes in PLC-γ1 critical for its activation^47, 48^. We speculate that the closed circular tetrameric formation mediated by the SLP-76-PLC-γ1 interaction would regulate the frequency and/or duration of SLP-76-bound Itk recruitment to the PLC-γ1 regulatory domain. Moreover, we previously demonstrated that replacement of PLC-γ1 in the tetramer by PLC-γ1 that had been phosphorylated at Y783 maximizes PLC-γ1 hydrolysis^6^. We thus propose that formation of the closed circular tetramer and phosphorylation of PLC-γ1 at Y783 co-operate to induce conformational changes necessary to optimize enzyme activation.

Replacement of SLP-76^WT^ with the SLP-76^HA^ protein led to a decrease in the number of post-positive selection DP3 thymocytes and post-negative selection medullary CD4 or CD8 SP most mature M2 thymocytes, accompanied by increased apoptosis. These results imply that the low affinity interaction between SLP-76 and PLC-γ1 acts as a checkpoint for T cell development. However, the degree of such decreases was very modest compared with that observed in LAT^G135D^ mice, suggesting that the SLP-76^HA^ mutation used in our study is less effective in enhancing TCR signal strength than the LAT^G135D^ mutation which accelerates the kinetics and degree of LAT phosphorylation at Y136^39^. We also noted a slight increase in the frequency of T cells with a memory phenotype (T_MP_) in the periphery of SLP-76^4Cre-mE^^8^ mice, but none of them showed any signs of autoimmunity as monitored for 1.5 years. In contrast, LAT^G135D^ mice spontaneously develop autoimmune diseases at 1 year of age, accompanied by a marked increase in dysfunctional Treg cells that fail to suppress the activation of T_MP_ cells, thus breaking peripheral tolerance^39^. However, the SLP-76^HA^ mutation barely affected the homoestasis and functional property of the Treg compartment, which would be capable of preventing the aberrant activation of autoreactive T_MP_ cells in SLP-76^4Cre-mE^^8^ mice.

We employed TCR transgenic systems to evaluate the effect of the SLP-76^HA^ mutation on the responsiveness of antigen-specific T cells. We showed that the SLP-76^HA^ mutation rendered OT-I CD8^+^ T cells more responsive to OVA APLs with lower affinities but not to a WT N4 peptide *in vitro*. However, the degree of enhanced TCR signal strength by the SLP-76^HA^ mutation was much smaller than that induced by the CD3ζ 6F(i) or LAT^G135D^ mutations, as judged by the EC_50_ of CD69 expression on OT-I cells stimulated with Q4H7 or an equivalent OVA APL^39, 54^. The latter two mutations lead to a substantial increase by over 10-fold and 100-fold, respectively, whereas the SLP-76^HA^ mutation resulted in only a 2-fold increase. Nonetheless, our observations that the SLP-76^HA^ accelerated the division of donor OT-I cells in response to Q4H7 but led to increased apoptosis in those cells with massive division *in vivo* highlight the biological relevance of the weak SLP-76-PLC-γ1 interaction in optimizing the responsiveness of peripheral T cells by fine-tuning TCR signal strength.

The potency of TCR signaling has been demonstrated as a key determinant underlying the fate decision of CD8^+^ T cells toward T_CM_ and T_EM_ cells^34, 35, 36, 37, 38, 39^. Using a P14 TCR transgenic and endogenous TCR systems, we showed that the SLP-76^HA^ mutation markedly reduced the generation of T_CM_ cells in the memory phase upon acute LCMV_Arm_ infection but had very little effect on the bifurcation into MPEC in the expansion phase. In contrast, the LAT^G135D^ mutation restrained OT-I cells from skewing toward MPEC at 7 days post-infection with *Listeria monocytogenes* expressing a WT OVA peptide or a low affinity V4 OVA APL^39^. These results indicate that the SLP-76^HA^ mutation is less potent than the LAT^G135D^ mutation at enhancing TCR signal strength in LCMV-reactive CD8^+^ T cells during the expansion phase when the virus-derived antigens are abundant, thus failing to restrain the fate decision toward MPEC. Notably, dampening TCR signal strength caused by the conditional SLP-76 deletion during the contraction phase enhanced LCMV-specific T_CM_ cells in the memory phase^35^. This leads us to speculate that the SLP-76^HA^ mutation would enhance TCR signal strength in LCMV_Arm_-primed CD8^+^ T cells that continuously recognize limited amounts of LCMV-derived peptide/MHC I complexes during the contraction phase, thus restraining the polarization toward T_CM_ cells.

Higher potency of TCR-mediated signaling has been demonstrated to favor the fate decision of naïve CD4^+^ T cells toward Tfh cells over other effector Th subsets^44, 45^. Using an OT-II CD4^+^ T cell adoptive transfer system, we showed that the SLP-76^HA^ mutation did not affect the differentiation of OT-II donor cells into pre-Tfh cells, although there was a slight reduction in the absolute number of donor-derived non-Tfh and pre-Tfh cells. These results indicate that the degree of enhanced TCR signal strength by the SLP-76^HA^ mutation is insufficient to force antigen-primed OT-II cells to undertake the Tfh differentiation initiation program^43^. Full polarization into mature Tfh cells requires a stable and prolonged interaction of pre-Tfh cells with B cells presenting a cognate antigen, where pre-Tfh cells tend to receive stronger TCR-mediated signaling^43^. We noted that the SLP-76^HA^ mutation led to a marked decrease in the number of mature Tfh cells, accompanied by increased death of CXCR5^+^ OT-II donor cells. Thus, the low affinity interaction of SLP-76 with PLC-γ1 fine-tunes TCR signal strength to prevent excessive death of pre-Tfh cells in the process of full maturation into functional Tfh cells for the maintenance of normal humoral immune responses.

There has been much attention recently on the central role of LAT phosphorylation on the Y132 residue in determining the response of TCR signaling^39, 55^. The kinetics of phosphorylation at this site, as regulated by the identity of the preceding glycine residue, is slower than the phosphorylation of the other three phosphorylated LAT residues. This slower phosphorylation acts as a proof-reading step necessary for defining the ultimate response by the T cell. Moreover, six tyrosine residues on immunoreceptor tyrosine-based activation motifs within the CD3ζ chains play a critical role in ligand discrimination by the TCR recognizing antigenic peptides with lower affinities in the context of MHC^54^. Our observations using the SLP-76^HA^ mutant shed light on the importance of the SLP-76 PRR_185-200_ in optimizing the activation of T cells. As the LAT-Gads-SLP-76-PLC-γ1 tetramer physiologically forms by the cooperative interaction of the four proteins following engagement of the TCR/CD3 complex and LAT phosphorylation, the SLP-76-PLC-γ1 interaction would serve as an additional regulatory kinetic proof-reading mechanism determining optimal TCR signaling.

Our present study, combining biophysical analyses, biochemical reconstitution, and use of a conditional knock-in mouse, indicate that it is the formation of the LAT-Gads-SLP-76-PLC-γ1 circular tetramer that is central to appropriate PLC-γ1 activation. Recruitment of PLC-γ1 to phosphorylated LAT associated with Gads and the SLP-76^NA^ mutant, which is unable to interact with PLC-γ1, was insufficient for full PLC-γ1 activation. Use of the SLP-76^HA^ mutant enhancing the affinity for PLC-γ1 by 2-3-fold over SLP-76^WT^ led to a slight but significant increase in PLC-γ1 activity. However, this small increase in the affinity of the SLP-76-PLC-γ1 interaction resulted in impaired thymocyte development and peripheral T cell responses *in vivo*. These results and the fact that the amino acid sequences of the relevant interacting sites of PLC-γ1 and SLP-76 are conserved lead to the conclusion that the affinities of protein-protein interaction defined in the tetramer formation, especially the weak SLP-76-PLC-γ1 interaction, are critical to the controlled activation of PLC-γ1. This weak interaction thus fine-tunes TCR signal strength to generate appropriate T cell-mediated immune responses for all vertebrates.

## Materials and Methods

### Protein Purification

Recombinant proteins were expressed in bacteria or insect cells and were purified using one or more of the following: Ni-affinity chromatography (5 mL HisTrap HP column, GE Healthcare); GST-affinity chromatography (Glutathione resin, GenScript); HaloTag-affinity chromatography (HaloTag Protein Purification Resins, Promega); Anion exchange column chromatography (ResourceQ, Cytiva Life Sciences); Size-exclusion column chromatography (SEC) (Superdex-200 increase 10/300 GL column, Cytiva Life Sciences). The concentrations of each protein were determined spectrophotometrically. The detailed purification procedures for PLC-γ1, Gads, SLP-76 and LAT were described in Manna et al.^3^ and Wada et al.^6^

The cDNA fragments of Halotagged PLC-γ1 and dual-tagged (His_6_– and Halo-) Gads were inserted into the pDest vector; each protein was expressed in Sf9 insect cells and harvested (Leidos Biomedical Research, Inc.). The protein lysates were prepared by French press and centrifugation. HaloTag-affinity resin and Ni-affinity column were used for PLC-γ1 and Gads purification respectively. The HaloTag on PLC-γ1 was cleaved away, but the HaloTag on Gads remained to enhance protein solubility. After the affinity purifications, these were further purified by SEC.

The SLP-76 cDNA (103-258aa with Y113F and Y173F mutation) linked to a His_6_-GST-TEV protease cleavage sequence at N-terminus was inserted into pTE28a vector. The protein was expressed in Rosetta 2(DE3)pLysS cells (Novagen). The protein lysate prepared by French press and centrifugation was incubated with glutathione resin (GenScript) for 2h at 4°C. After washing the resin, the SLP-76 protein was cleaved off from the resin with TEV protease. The liberated SLP-76 protein and the His_6_-TEV protease were separated by Ni-affinity chromatography with a shallow imidazole gradient. The eluted SLP-76 protein was further purified by SEC.

The cDNA fragment of LAT (48-233aa) linked to His_6_– and MBP-tag sequences at the N-terminus was inserted into the pET28a vector. The protein was expressed in Rosetta2(DE3)pLysS cells. Multi-tagged LAT protein was purified from the protein lysate by Ni-affinity column and ion-exchange column chromatography (IEX). The His_6_– and MBP-tag at the N-terminus were removed using TEV protease. The cleaved LAT protein was purified by the same IEX again and then treated with *in vitro* phosphorylation. The phosphorylated LAT was passed through a final IEX and SEC. Phosphorylation was confirmed by mass spectrometry.

The expression and purification of Itk protein: His_6_-3C-Halo-tev-ITK(C96E/T110I)-FLAG were carried out following a previously established protocol^56^.

### Enzymatic assay for PIP2 cleavage by PLC-γ1

The detailed experimental procedures were described in Wada et al.^6^ Large Unilamellar Vesicles (LUVs) provided lipid bilayer surfaces as a surrogate of the cellular membrane environment and also contained PIP2, a substrate for the PLC-γ1 enzymatic reaction. The LUVs were prepared using a Mini extruder kit (Avanti Polar Lipids) and comprised of phosphatidylcholine (PC), phosphatidylethanolamine (PE), phosphatidylserine (PS), phosphatidylinositol 4,5-bisphosphate (PIP2), cholesterol, and 1,2-dioleoyl-sn-glycero-3-[(N-(5-amino-1-carboxypentyl) iminodiacetic acid) succinyl]; DGS-NTA(Ni). At the enzymatic reaction step, the proteins were mixed and incubated at 30°C, and the reaction was initiated by adding the LUVs. The diameter of the LUV was 0.4 µm, and protein concentrations were 75 nM of PLC-γ1, 375 nM of LAT, 562 nM of Gads and SLP-76. The reaction was stopped by flash freezing in liquid nitrogen. After removing the LUVs and proteins through a Captiva EMR Lipid (Agilent) and a Macro Spin Column C18 (Harvard Apparatus), the IP3 in the elution was quantified by HPLC analysis as described in Mayr et al.^57, 58^ The quantification analysis was performed by Image J.

### Isothermal titration calorimetry for measuring the affinity of protein interaction

The reference experimental procedures were described in Manna et al.^3^ and Houtman et al.^5^ ITC measurements were performed using a VP-ITC calorimeter or PEAQ-ITC (Malvern Panalytical). Titrations were performed in 20 mM Hepes (pH 7.4), 100 mM NaCl, with 0.1 mM DTT at 5°C. The initial protein concentration of SLP-76 in the cell and that of PLC-γ1 or Itk in the syringe were 4 µM and 40 µM, respectively. The injections of 8 µL were repeated 29 times in VP-ITC. The injections of 2 µL were repeated 19 times in PEAQ-ITC. Heat changes in each injection were fitted with the single binding site mechanism with 1:1 stoichiometry. The values for the stoichiometry of binding and thermodynamic constants were determined using the software package provided by Malvern Panalytical.

### Sequence alignment analysis of proline-rich region in SLP-76 and SH3 domain in PLC-γ1

Nineteen sequences of SLP-76 orthologs in jawed vertebrates from cartilaginous fish to primates, were selected. The sequences were searched by UniProt and Ensembl database, and the alignment analysis was performed using Protein BLAST.

### Mice

All mice used in the present study were on a C57BL/6 background. C57BL/6J (B6/J) mice (Strain #: 000664), B6 CD45.1 mice (B6.SJL-*Ptprc^a^Pepc^b^*/BoyJ; Strain #: 002014)^33^, CD4^Cre^ mice (B6.Cg-Tg(Cd4-cre)1Cwi/BfluJ; Strain #: 022071)^17^, OT-I mice (C57BL/6-Tg(TcraTcrb)1100Mjb/J; Strain #: 003831)^31^, OT-II mice (B6.Cg-Tg(TcraTcrb)425Cbn/J; Strain #: 004194)^46^, and R26R-EYFP mice (B6.129X1-Gt(ROSA)26Sor^tm1^(EYFP)*^Cos^*/J; Strain #: 006148)^16^ were purchased from The Jackson Laboratory. C57BL/6Ncr (B6/Ncr) mice used as a source of fertilized one-cell embryos for microinjection were purchased from Charles River Laboratories. P14 mice (C57BL/6Ai-[Tg]TCR P14 LCMV N10) were purchased from Taconic^40^. hCD2^Cre^ mice (C57BL/6-Tg(CD2-cre)1Lov/J) were provided by Dr. Paul. E. Love at NICHD, NIH^30^. Mice were housed in specific pathogen-free facilities and used between 8 and 20 weeks of age. Both female and male mice were used except for adoptive T cell transfer experiments where only females were used as a source of donor T cells. Animal procedures were approved by the NCI Animal Care and Use Committee.

### Generation of Lcp2^L^^8:21L-mE^^8^ mice

We utilized the CRISPR/Cas9 system to generate Lcp2^L8:21L-mE8^ mice. Briefly, four single guide (sg) RNAs were designed using sgRNA Scorer 2.0 (PMID: 28146356) and subsequently tested for *in vitro* cutting activity measured by high throughput sequencing of which sgRNA-1: AGAGCCGTTTAATTCCATCC and sgRNA-2: AACCACTGAACAGAACGTGA were chosen for subsequent mouse generation. To generate the donor/targeting vector, a double stranded plasmid DNA was first cloned using a combination of synthesized DNA fragments (Twist Bioscience) and isothermal assembly (PMID: 19363495) into the pGMC00018 backbone (Addgene #195320). This plasmid consisted of WT SLP-76 cDNA spanning exons 8 through 21 followed by a poly A sequence, which was flanked by LoxP sequences. The second LoxP sequence was followed by the mutated exon 8 encoding the SLP-76^HA^. Upon Sanger sequence validation of this construct, a long single stranded DNA (ssDNA) was then generated using the Guide-it Long ssDNA Production system (Takara Bio). Finally, Cas9 protein, chemically modified versions of the above listed sgRNAs (Synthego) targeting the upstream and downstream introns of exon 8, and the long ssDNA fragment were then microinjected into fertilized one-cell embryos of B6/Ncr mice. After microinjection the embryos were transferred into the oviduct of pseudo-pregnant B6D2F1 females. Offspring from the recipient females were genotyped to identify a founder mouse carrying the successfully recombineered *Lcp2^L^*^8^*^:21L-mE^*^8^ allele by PCR using two sets of primers, as depicted in Fig. S2A. GT-1F: CATGCACACGGATCATGTTGC, GT-1R: TGCTATACGAAGTTATAGAGCCG, GT-2F: TGGGTCGTTTGTTCATAACTTCG, and GT-2R: ACTGGTGTGAAGTAGCTGCC. The GT-1R primer spanned the upstream intron of exon 8 and the first LoxP, and the GT-2F primer spanned the second LoxP and the downstream intron of exon 8. The founder mouse was backcrossed onto B6/J mice at least 8 generations prior to further breeding as described below.

### Mouse breeding

To evaluate the effect of the SLP-76^HA^ mutation on thymocyte development and on peripheral T cell activation and differentiation, Lcp2^L8:21L-mE8^ mice were bred onto CD4^Cre^ mice and hCD2^Cre^ mice, respectively. The Lcp2^L8:21L-mE8^ mice carrying a CD4^Cre^ or hCD2^Cre^ transgene were further mated with R26R-EYFP mice to induce YFP expression in cells with Cre activity, thus marking the mutated SLP-76-expresssing cells. R26R-EYFP mice carrying a CD4^Cre^ or hCD2^Cre^ transgene were used as a control. To assess the effect of the SLP-76^HA^ mutation on antigen-specific T cell responses, Lcp2^L8:21L-mE8^ x R26R-EYFP x hCD2^Cre^ mice were further crossed onto OT-I, P14, or OT-II mice. R26R-EYFP x hCD2^Cre^ mice carrying a corresponding TCR transgene were used as a control. In the resultant mouse lines generated by the genetic cross, Lcp2^L8:21L-mE8^ and R26R-EYFP alleles were maintained homozygous, but the Cre and TCR transgenes were maintained hemizygous. (CD45.1 x CD45.2) F1 mice were generated by breeding B6 CD45.1 mice onto B6/J mice and used as recipients for adoptive T cell transfer experiments.

### Antibodies

Biotinylated anti-γδ TCR (GL3), BUV395 anti-CD4 (RM4-5), BUV737 anti-CD45.2 (104), BUV737 anti-TCRβ (H57-597), BUV805 anti-CD44 (IM7), BV510 anti-CD19 (1D3), BV510 anti-γδ TCR (GL3), BV650 anti-CD103 (2E7), BV650 anti-CD122 (TM-β1), BV650 anti-H-2K^b^ (AF6-88.5), BV711 anti-CD24 (M1/69), BV711 anti-PSGL-1 (2PH1), BV786 anti-Vα2 TCR (B20.1), RB705 anti-TCF1 (S33-966), RB780 anti-T-bet (O4-46), PE anti-SAP (1A9), PE-CF594 anti-Blimp-1 (5E7), PE-Cy7 anti-CD45.1 (A20), Alexa Flour 647 anti-Bcl-6 (K112-91), and R718 anti-CD3 (17A2) were purchased from BD Biosciences. Biotinylated antibodies to CD3 (17A2), CD4 (RM4-5), CD8a (53-6.7), CD11b (M1/70), CD11c (N418), CD25 (PC61), CD49b (DX5), CD69 (H1.2F3), CD90.2 (30-H12), NK-1.1 (PK136), ICOS (C398.4A), PD-1 (29F.1A12), I-A/I-E (M5/114.15.2), and TER-119 (TER-119), BV421 anti-CCR7 (4B12), BV421 anti-CXCR5 (L138D7), BV421 anti-PD-1 (29F.1A12), BV421 anti-KLRG1 (2F1/KLRG1), BV510 anti-CXCR3 (CXCR3-173), BV510 anti-I-A/I-E (M5/114.15.2), BV510 anti-NK-1.1 (PK136), BV605 anti-CD62L (MEL-14), BV711 CD25 (PC61), BV785 anti-I-A/I-E (M5/114.15.2), BV785 anti-Ly-6C (HK1.4), Alexa Flour 488 anti-GFP (FM264G; This antibody cross-reacts with YFP), PerCP-Cy5.5 anti-CD5 (53-7.3), PE anti-PD-1 (29F.1A12), PE-Dazzle 594 anti-CXCR5 (L138D7), PE-Dazzle 594 anti-PD-1 (29F.1A12), PE-Cy5 anti-CD127 (A7R34), PE-Cy7 anti-CD25 (PC61), APC anti-CD69 (H1.2F3), APC anti-Helios (22F6), and purified anti-CD16/CD32 (93) were purchased from BioLegend. Biotinylated antibodies to CD16/CD32 (93) and CD19 (1D3), PE-eFluor 610 anti-Foxp3 (FJK-16s), and APC-eFluor 780 anti-CD8a (53-6.7) were purchased from eBiosciences. PE anti-Cleaved Caspase-3 (D3E9) and APC anti-TOX (REA473) were purchased from Cell Signaling Technology and Miltenyi Biotec, respectively.

### Isolation of T cells and T cell-depleted spleen cells

A single cell suspension prepared from whole LNs (axillary, brachial, superficial/deep cervical, inguinal, and mesenteric) and spleens of T cell donor mice was treated with ACK lysis buffer (KD Medicals) to deplete red blood cells (RBC). The remaining cells were suspended in 1X PBS (KD Medicals) containing 2% fetal bovine serum (FBS; Gibco) and incubated with biotinylated antibodies to CD4 (for CD8^+^ T cell isolation), CD8a (for CD4^+^ T cell isolation), CD19, I-A/I-E, CD16/CD32, NK-1.1, CD25, γδ TCR, CD11b, CD11c, CD49b, CD69, ICOS, PD-1, TER-119. After washing extensively, the antibody-bound cells were incubated with Streptavidin Microbeads (Miltenyi) and loaded onto LS Columns (Miltenyi), following the manufacturer’s instruction. The magnetically unbound cells were used for an *in vitro* T cell activation assay and for adoptive transfer to assess the responsiveness of donor T cells *in vivo*. For isolation of T cell-depleted spleen (TdS) cells, a single cell suspension was prepared from spleens of (CD45.1 x CD45.2) F1 mice. Most of the procedures were the same as the T cell isolation except that the remaining cells after RBC lysis were incubated with biotinylated antibodies to CD3, γδ TCR, and CD90.2. The magnetically unbound cells were used as antigen-presenting cells for *in vitro* T cell activation assay.

### In vitro OT-I CD8^+^ T cell activation

TdS cells from (CD45.1 x CD45.2) F1 mice were loaded with varying concentrations of WT OVA peptide N4 (SII<u>N</u>FEKL; Anaspec), Q4H7 (SII<u>Q</u>FE<u>H</u>L; Anaspec), or G4 (SII<u>G</u>FEKL; Anaspec) in 200 µL of RPMI 1640 (Gibco) containing 5% FBS, 1 mM Sodium Pyruvate (Gibco), 1X MEM Non-Essential Amino Acids (Gibco), 2 mM L-Glutamine (Gibco), and 100 U/mL Penicillin-Streptomycin (Gibco) (thereafter cRPMI) on a 96-well round bottom cell culture plate (Coster) at 37℃ for 3 hours in a humidified CO_2_ incubator. After peptide loading, TdS cells were washed 3 times with 1X PBS. CD8^+^ T cells isolated from WT OT-I CD45.1 and SLP-76^2Cre-mE8^ OT-I CD45.2 mice were mixed at a 1:1 ratio and stimulated with peptide-loaded TdS cells for 20 hours on a 96-well round bottom cell culture plate. Each well contained 5 x 10^4^ of WT CD45.1 OT-I cells, 5 x 10^4^ of SLP-76^2Cre-mE8^ CD45.2 OT-I cells, and 1 x 10^5^ of peptide-loaded (CD45.1 x CD45.2) F1 TdS cells in 200 µL of cRPMI.

### Adoptive T cell transfer and immunization

CD8^+^ T cells isolated from WT OT-I CD45.1 and SLP-76^2Cre-mE8^ OT-I CD45.2 mice were mixed at a 1:1 ratio and labeled with 5 µM CellTrace Violet (Invitrogen) following the manufacturer’s instruction. After CTV labeling and extensive washing with 1X PBS, cells were suspended in 1X PBS at 1 x 10^6^ cells/mL. (CD45.1 x CD45.2) F1 mice were injected with 100 µL of cell suspension (containing 5 x 10^4^ OT-I cells of each genotype) intravenously via retro-orbital sinus. One day later, recipient mice were immunized with 10 nmol of Q4H7 emulsified in complete Freund’s adjuvant (CFA; Sigma) subcutaneously via left and right lateral. Donor OT-I cells in skin dLNs (axillary, brachial, and inguinal) were analyzed at 3 days post-immunization by flow cytometry.

To assess the responsiveness of CD4^+^ T cells *in vivo*, 1:1-mixed OT-II cells isolated from WT OT-II CD45.1 and SLP-76^2Cre-mE8^ OT-II CD45.2 mice were suspended in 1X PBS at 1 x 10^6^ cells/mL and injected into (CD45.1 x CD45.2) F1 mice with a volume of 100 µL containing 5 x 10^4^ OT-II cells of each genotype. One day later, recipient mice were immunized with 50 µg of EndoFit OVA (Invivogen) in CFA subcutaneously via left and right lateral. Donor OT-II cells in skin dLNs were analyzed at 6, 8, and 10 days post-immunization by flow cytometry.

### LCMV infection

CD8^+^ T cells isolated from WT P14 CD45.1 and SLP-76^2Cre-mE8^ P14 CD45.2 mice were mixed at a 1:1 ratio and suspended in 1X PBS at 1 x 10^5^ cells/mL. (CD45.1 x CD45.2) F1 mice were injected with 100 µL of cell suspension (containing 5 x 10^3^ P14 cells of each genotype) in PBS intravenously via retro-orbital sinus. One day later, recipient mice were infected intraperitoneally with 5 x 10^4^ plaque-forming units of LCMV_Arm_. Donor P14 cells in the spleens of infected recipients were analyzed at 7 and 28 days post-infection by flow cytometry. In some experiments, WT and SLP-76^2Cre-mE8^ mice bearing endogenous TCR repertoire were infected with the same dose of LCMV_Arm_ to assess the virus-specific CD8^+^ T cells at 30 days post-infection.

### Cell preparation and staining for flow cytometry analysis

For characterization of thymocytes, single cell suspensions prepared from thymi of WT and SLP-76^4Cre-mE8^ mice at 8 – 12 weeks of age. Cells were stained with Blue Fluorescent Reactive Dye (Invitrogen) for dead cell exclusion, followed by surface staining with BUV395 anti-CD4, BUV737 anti-TCRβ, BV805 anti-CD44, BV421 anti-CCR7, BV605 anti-CD62L, BV650 anti-H-2K^b^, BV711 anti-CD24, BV785 anti-Ly-6C, PerCP-Cy5.5 anti-CD5, PE-Cy5 anti-CD127, PE-Cy7 anti-CD25, APC anti-CD69, APC-eFluor 780 anti-CD8a in the presence of purified anti-CD16/CD32 to prevent FcR-mediated non-specific binding of the antibodies to surface molecules. The stained thymocytes were fixed/permeabilized first with Cytofix/Cytoperm Solution (BD Biosciences) for 20 minutes and then with Foxp3/Transcription Factor Fixation/Permeabilization Solution (eBiosciences) for 30 minutes, followed by intracellular staining with Alexa Flour 488 anti-GFP, PE anti-Cleaved Caspase-3, PE-eFlour 610 anti-Foxp3, R718 anti-CD3. Pre-fixation/permeabilization with Cytofix/Cytoperm Solution prevented the loss of YFP signal caused by Foxp3/Transcription Factor Fixation/Permeabilization Solution.

To characterize peripheral T cells from WT and SLP-76^4Cre-mE8^ mice at 8 – 12 weeks of age, pooled whole LNs (axillary, brachial, superficial/deep cervical, inguinal, and mesenteric) were used for preparation of single cell suspension. Following Blue Fluorescent Reactive Dye staining, cells were stained with BUV395 anti-CD4, BUV737 anti-TCRβ, BV805 anti-CD44, BV421 anti-CXCR5, BV605 anti-CD62L, BV650 anti-CD122, BV711 anti-PSGL-1, BV785 anti-Ly-6C, PerCP-Cy5.5 anti-CD5, PE anti-PD-1, PE-Cy5 anti-CD127, PE-Cy7 anti-CD25, APC-eFluor 780 anti-CD8a in the presence of purified anti-CD16/CD32. After surface staining, cells were fixed/permeabilized in the same way as thymocytes and stained intracellularly with Alexa Flour 488 anti-GFP, PE-eFluor 610 anti-Foxp3, APC anti-Helios, R718 anti-CD3.

To analyze *in vitro* activated OT-I cells, cultured cells were stained with Blue Fluorescent Reactive Dye and then with BUV395 anti-CD4, BUV805 anti-CD44, BV605 anti-CD62L, BV711 CD25, BV786 anti-Vα2 TCR, PE anti-PD-1, PE-Cy5 anti-CD127, APC anti-CD69, and APC-eFluor 780 anti-CD8a in the presence of purified anti-CD16/CD32. After surface staining, cells were treated with Cytofix/Cytoperm Solution and stained intracellularly with BUV737 anti-CD45.2, Alexa Flour 488 anti-GFP, PE-Cy7 anti-CD45.1, and R718 anti-CD3.

To analyze CTV-labeled OT-I cells primed with Q4H7 *in vivo*, skin dLN cells were stained with Blue Fluorescent Reactive Dye, followed by surface staining with BUV395 anti-CD4, BUV805 anti-CD44, BV510 anti-I-A/I-E, BV510 anti-CD19, BV510 anti-NK-1.1, BV510 anti-γδ TCR, BV605 anti-CD62L, BV650 anti-CD103, BV711 anti-CD25, BV786 anti-Vα2 TCR, PerCP-Cy5.5 anti-CD5, PE-Dazzle 594 anti-PD-1, PE-Cy5 anti-CD127, APC anti-CD69, and APC-eFluor 780 anti-CD8a in the presence of purified anti-CD16/CD32. After surface staining, cells were fixed/permeabilized in the same way as *in vitro* activated OT-I cells and stained intracellularly with BUV737 anti-CD45.2, Alexa Flour 488 anti-GFP, PE anti-Cleaved Caspase-3, PE-Cy7 anti-CD45.1, and R718 anti-CD3.

For characterization of P14 donor cells in LCMV_Arm_-infected recipient mice, spleens were excised for preparation of single cell suspension. Following Blue Fluorescent Reactive Dye staining, cells were stained with BUV395 anti-CD4, BUV805 anti-CD44, BV421 anti-KLRG1, BV510 anti-CXCR3, BV605 anti-CD62L, BV650 anti-CD103, BV711 anti-CD25, BV785 anti-I-A/I-E, PE-anti-PD-1, PE-Cy5 anti-CD127, and APC-eFluor 780 anti-CD8a in the presence of purified anti-CD16/CD32. After surface staining, cells were fixed/permeabilized in the same way as thymocytes and stained intracellularly with BUV737 anti-CD45.2, Alexa Flour 488 anti-GFP, RB705 anti-TCF1, RB780 anti-T-bet, PE-CF594 anti-Blimp-1, PE-Cy7 anti-CD45.1, APC anti-TOX, and R718 anti-CD3. In some experiments, LCMV-reactive CD8^+^ T cells were detected using PE-conjugated LCMV_gp33-41_/H-2D^b^ tetramer (NIH Tetramer Core Facility) in the presence of 2 nM Dasatinib (Selleckchem) to prevent the downregulation of the tetramer-bound TCR. The tetramer staining was performed prior to staining for surface markers.

For characterization of OT-II donor cells in OVA/CFA-immunized recipient mice, skin dLN cells were stained with Blue Fluorescent Reactive Dye, followed by surface staining with BUV395 anti-CD4, BUV805 anti-CD44, BV421 anti-PD-1, BV510 anti-I-A/I-E, BV605 anti-CD62L, BV711 anti-PSGL-1, BV786 anti-Vα2 TCR, PE-Dazzle 594 anti-CXCR5, PE-Cy5 anti-CD127, and APC-eFluor 780 anti-CD8a in the presence of purified anti-CD16/CD32. After surface staining, cells were fixed/permeabilized in the same way as thymocytes and stained intracellularly with BUV737 anti-CD45.2, Alexa Flour 488 anti-GFP, RB705 anti-TCF1, RB780 anti-T-bet, PE anti-SAP, PE-Cy7 anti-CD45.1, Alexa Flour 647 anti-Bcl-6, and R718 anti-CD3.

Stained cells were run on the FACSymphony A5 (BD Biosciences), and the acquired flow cytometry data were analyzed with FlowJo 10 software (BD Biosciences). Gating strategies to analyze the populations of interest were shown in the Supplemental Figures.

### Statistical analyses

All statistical analyses were done with Prism 10 software (GraphPad Software). Error bars in graphs indicate average ± SD unless otherwise indicated. Comparisons were done by two-sided Student’s t-test, paired t-test, ordinary one-way ANOVA, or two-way ANOVA. For statistical comparisons, sample size was always greater than five and determined empirically based on pilot analyses. Animals were excluded from analyses only because of poor health status unrelated to the experiment.

## Acknowledgments

We thank Dr. Paul E. Love (NICHD, NIH) and the NIH Tetramer Core Facility for providing us with hCD2^Cre^ mice and PE-conjugated LCMV_gp33-41_/H-2D^b^ tetramer, respectively. We are grateful to the staff at NCI-Bethesda Building 37 and NCI-Frederick Animal Facilities for their excellent husbandry of our mouse colony. We also thank Dr. Paul A. Randazzo (LCMB, CCR, NCI, NIH) for his critical reading of the manuscript.

## Author contributions

L.E.S. conceptualized the study. H.Y., J.W., M.S.V., R.B., and L.E.S. designed the study. H.Y., J.W., E.N.S., N.D.C., M.E.L., and M.S.V. performed the experiments. W.L. performed mouse genotyping. U.M.R. prepared purified Itk for ITC assay. L.B., R.C., H.H., and P.A. helped generate Lcp2^L^^8:21L-mE8^ mice. D.B.M. prepared the stock of LCMV_Arm_. H.Y., J.W., E.N.S., N.D.C., and M.E.L. interpreted data and prepared figures. H.Y., J.W., and L.E.S. wrote the manuscript with comments from M.E.L., M.S.V., L.B., R.B., and P.A.R.

## Data and materials availability

All data required for the authors to draw the conclusions are present in the current study and available from the corresponding authors. Reagents used in the current study will be available upon request to the corresponding authors after completing a materials transfer agreement with NCI, NIH.

## Extended data figure legends

**Extended Data Figure 1:**
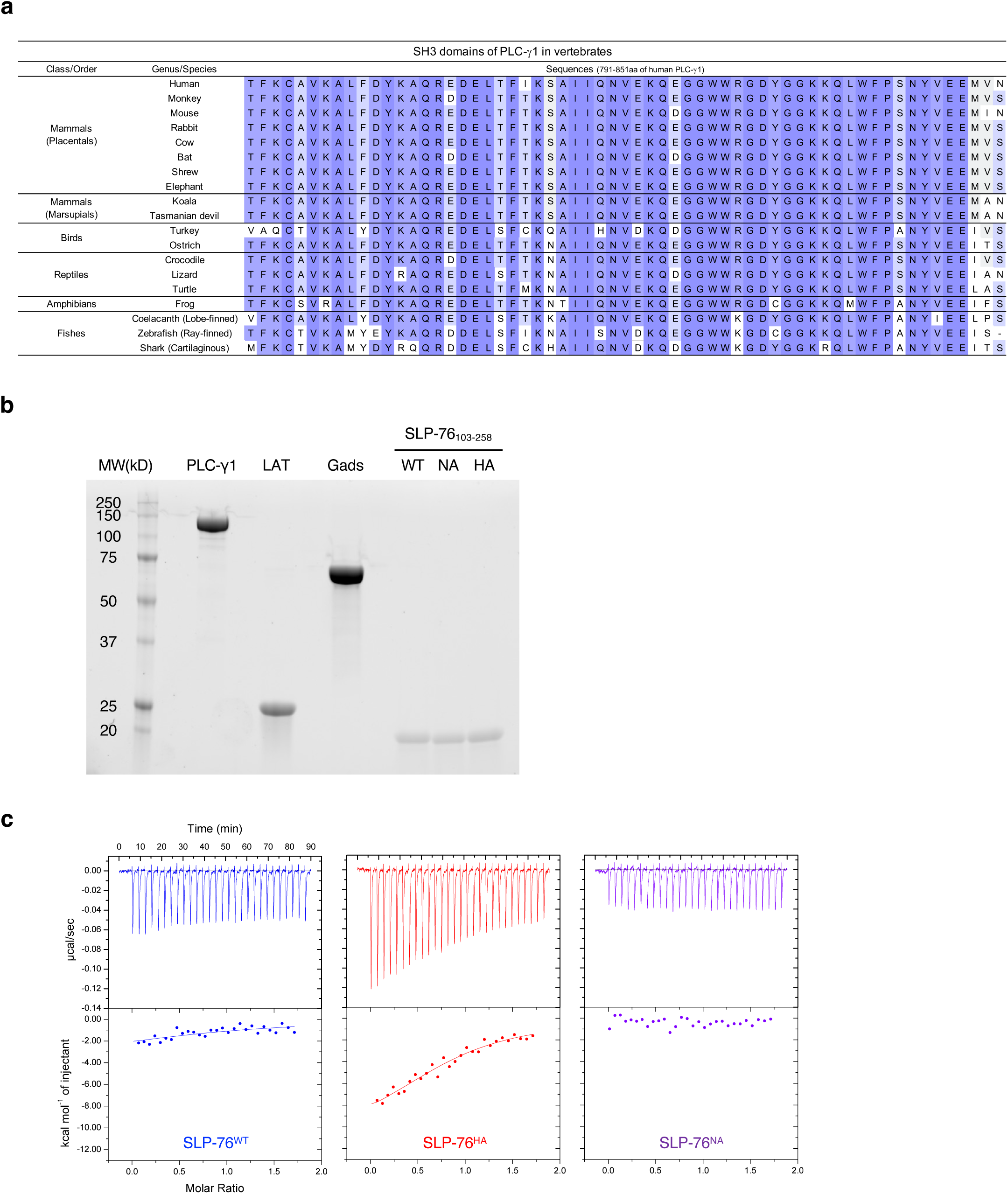
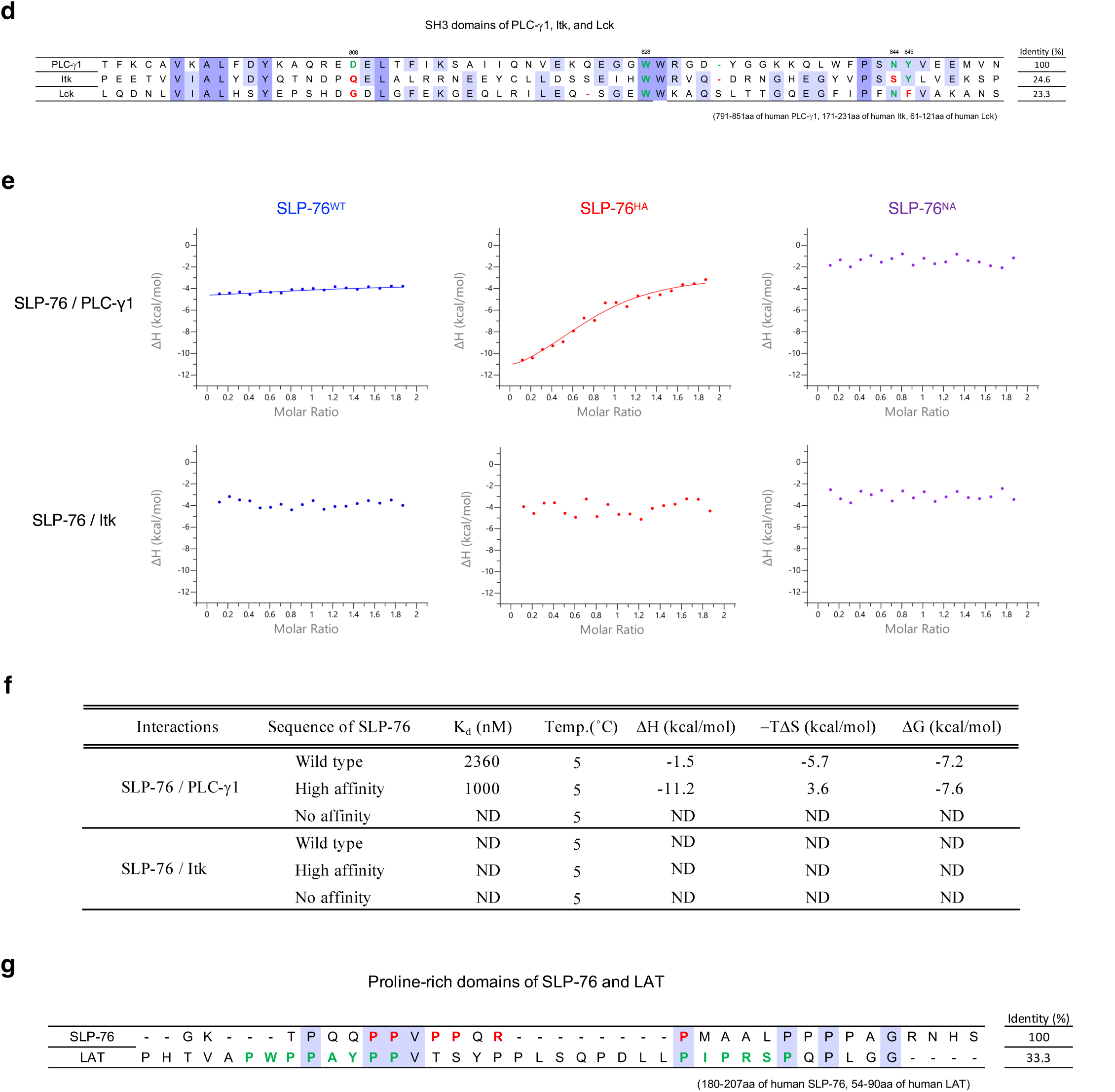
*In vitro* reconstitution studies of tetramer formation (corresponding to Figure 1). **a**, Sequence alignment of the SH3 domain of PLC-γ1 among 19 jawed vertebrates. **b**, Purified recombinant proteins used in the current study were resolved by SDS-PAGE and imaged with the Chemi-doc system. **c**, Spectrum of isothermal titration calorimetry for the interaction of SLP-76 peptides with PLC-γ1 measured by VP-ITC. **d**, Sequence alignment of the SH3 domains of PLC-γ1, Itk, and Lck. PLC-γ1 residues D808, W828, N844, and Y845, in green, are critical residues for binding SLP-76^8^. In both Itk and Lck, multiple residues in red corresponding to those critical residues in PLC-γ1 are mutated. **e**, Spectrum of isothermal titration calorimetry for the interaction of SLP-76 with PLC-γ1 or Itk, measured by PEAQ-ITC. **f**, Thermodynamic parameters of the interaction of SLP-76 with PLC-γ1 or Itk. **g**, Sequence alignment of the proline-rich regions of SLP-76 and LAT. In the SLP-76 sequence, prolines and arginine in red are critical residues for binding to PLC-γ1. In the LAT sequence, amino acids in green are critical residues for binding to Lck^14^.

**Extended Data Figure 2:**
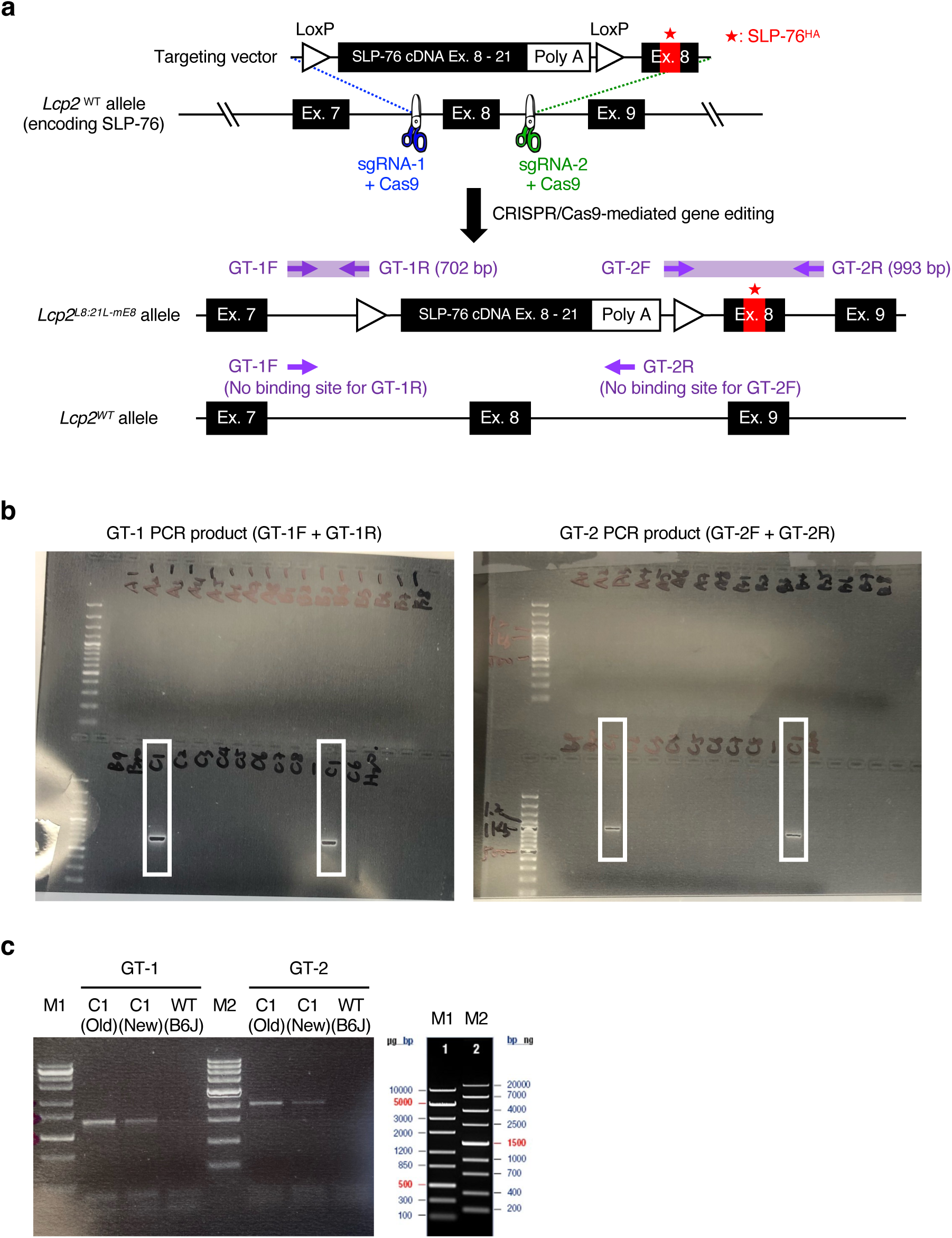
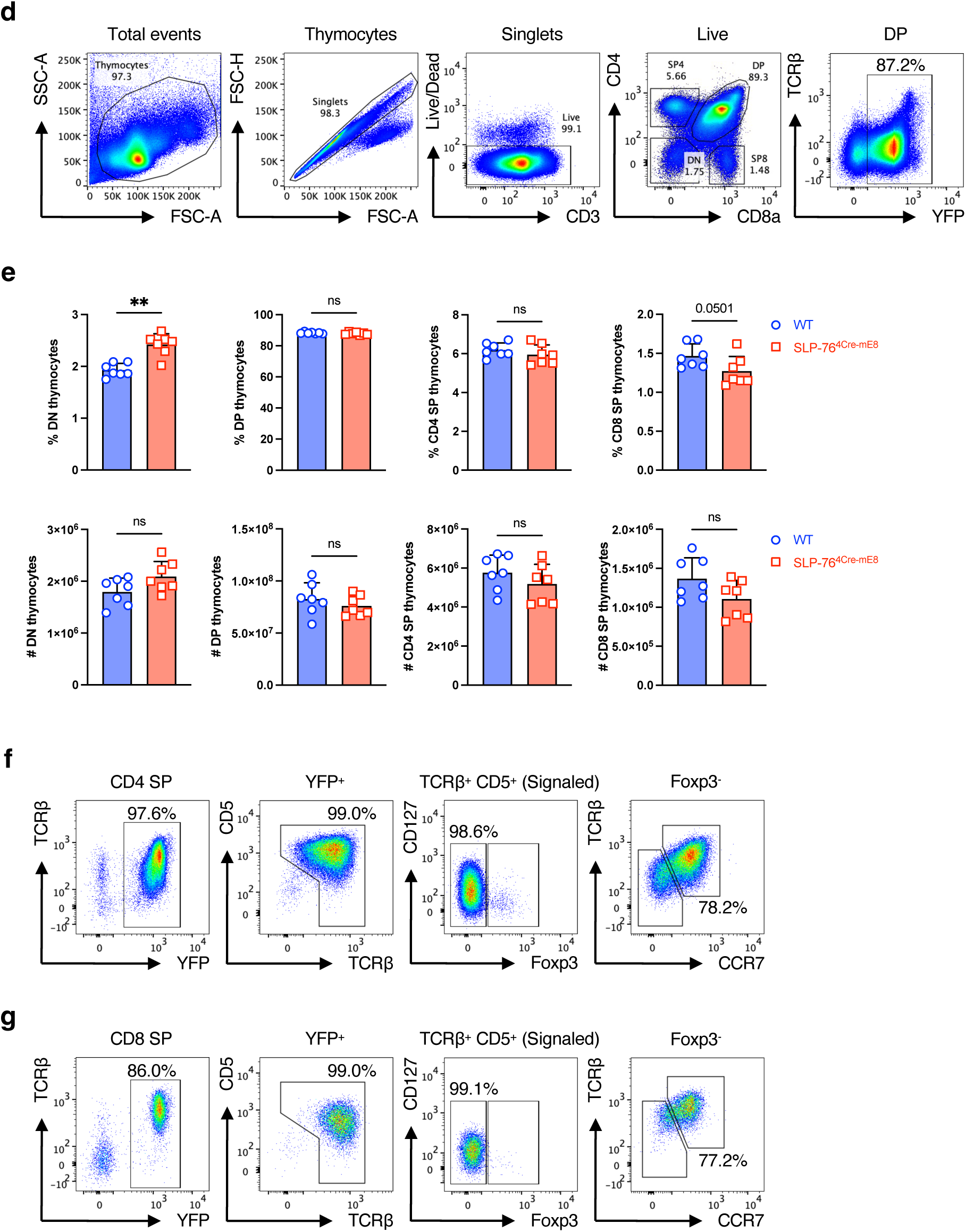
Generation of Lcp2^L8:21L-mE8^ mice and gating strategies for analysis of thymocytes (corresponding to Figure 2). **a**, Diagram of gene targeting strategy to generate an Lcp2^L8:21L-mE8^ mouse. **b**, Identification of a founder mouse carrying the recombineered *Lcp2^L^*^8^*^:21L-mE^*^8^ allele by PCR using two sets of primers, GT-1 and GT-2. **c**, Further confirmation of the founder mouse C1 by PCR using GT-1 and GT-2 primers. **d**, Gating strategy for analysis of YFP^+^ CD4/CD8 DP thymocytes. **e**, Statistical analysis for the frequency (upper row) and absolute number (lower row) of DN, DP, CD4 SP, and CD8 SP thymocytes. **f**,**g**, Gating strategy for analysis of YFP^+^ CD4 SP (**f**) and CD8 SP thymocytes (**g**). Statistics was determined by two-sided Student’s t-test. ns; not significant, **; p<0.01.

**Extended Data Figure 3:**
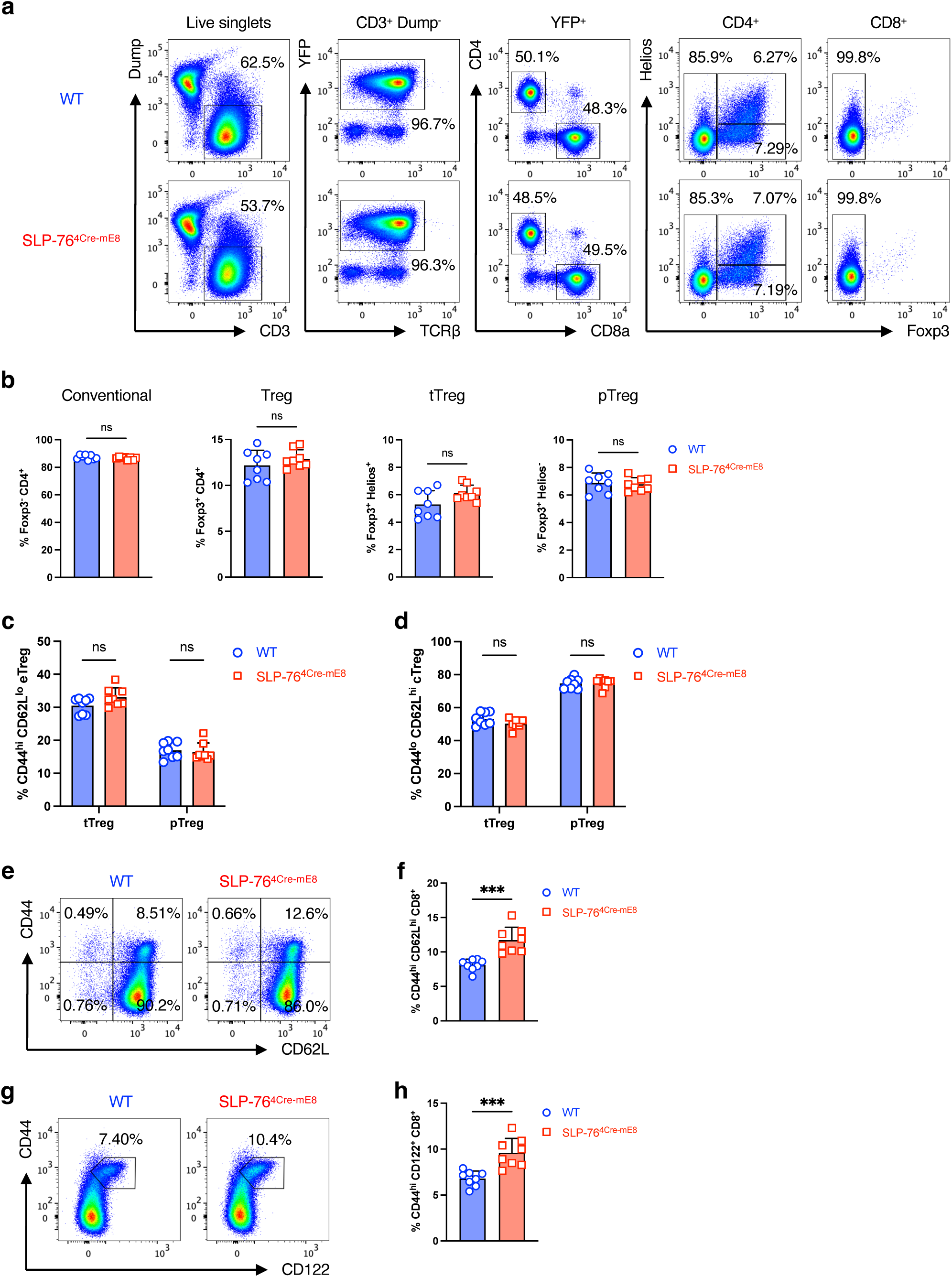
Gating strategies for analysis of peripheral T cells (corresponding to Figure 3). **a**, Gating strategy for analysis of YFP^+^CD4^+^ and YFP^+^CD8^+^ T cells in pooled whole LNs from WT (top) and SLP-76^4Cre-mE8^ (bottom) mice. Dump channel contains CD19, I-A^b^, NK-1.1, and γδ TCR. **b**, Statistical analysis of CD3^+^YFP^+^CD4^+^ T cells for the frequency of conventional, total Tregs, tTregs, and pTregs (n=8). **c**,**d**, Statistical analysis of tTreg and pTreg compartments for the frequency of eTregs (**c**) and cTregs (**d**) (n=8). **e**,**f**, Representative pseudocolor plot analysis of CD3^+^YFP^+^Foxp3^-^ conventional CD8^+^ T cells for CD62L and CD44 expressions (**e**) and statistical analysis for the frequency of CD44^high^CD62L^high^ CD8^+^ T cells corresponding to CD8^+^ T_MP_ cells (**f**) (n=8). **g**,**h**, Representative pseudocolor plot analysis of CD3^+^YFP^+^Foxp3^-^conventional CD8^+^ T cells for CD62L and CD122 expressions (**g**) and statistical analysis for the frequency of CD44^high^CD122^+^ CD8^+^ T cells corresponding to CD8^+^ T_MP_ cells (**h**) (n=8). Statistics was determined by two-sided Student’s t-test. ns; not significant, ***; p<0.001.

**Extended Data Figure 4:**
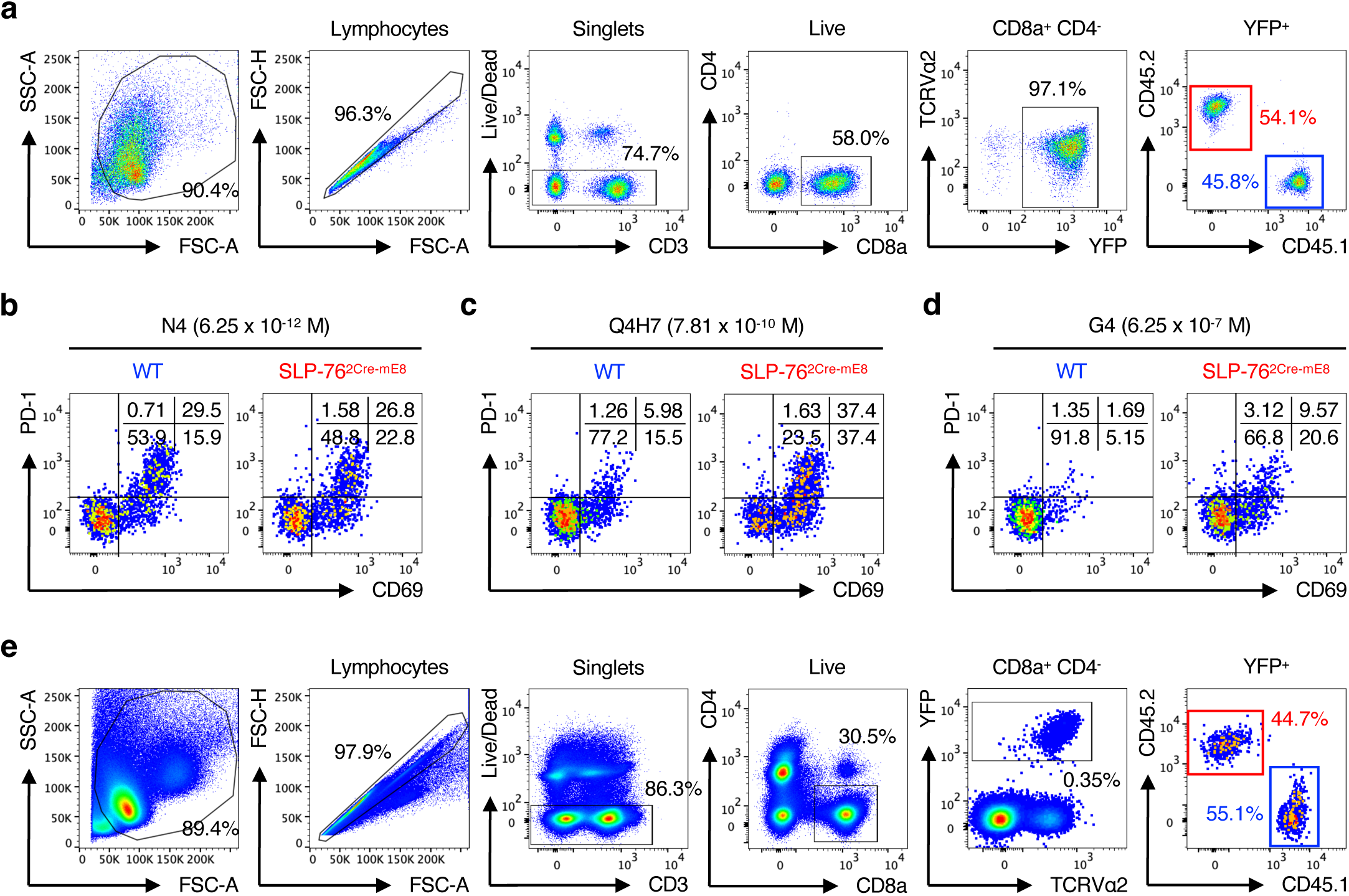
Gating strategy for analysis of WT and SLP-76^2Cre-mE8^ OT-I CD8^+^ T cells stimulated with peptide-loaded TdS cells, and representative pseudo plot analysis for CD69 and PD-1 expressions (corresponding to Figure 4). **a**, Gating strategy for analysis of OT-I CD8^+^ T cells stimulated with peptide-loaded TdS cells. **b**–**d**, Representative pseudocolor plot analysis of WT CD45.1 and SLP-76^2Cre-mE8^ CD45.2 OT-I CD8^+^ T cells for CD69 and PD-1 expressions upon *in vitro* stimulation with 6.25 x 10^-12^ M N4 (**b**), 7.81 x 10^-10^ M Q4H7 (**c**), or 6.25 x 10^-7^ M G4 (**d**). The numbers indicate the frequency of cells in each quadrant. **e**, Gating strategy for analysis of co-transferred WT CD45.1 and SLP-76^2Cre-mE8^ CD45.2 OT-I CD8^+^ T cells in skin dLNs of (CD45.1 x CD45.2) F1 recipient mice immunized with Q4H7 in CFA for 3 days.

**Extended Data Figure 5:**
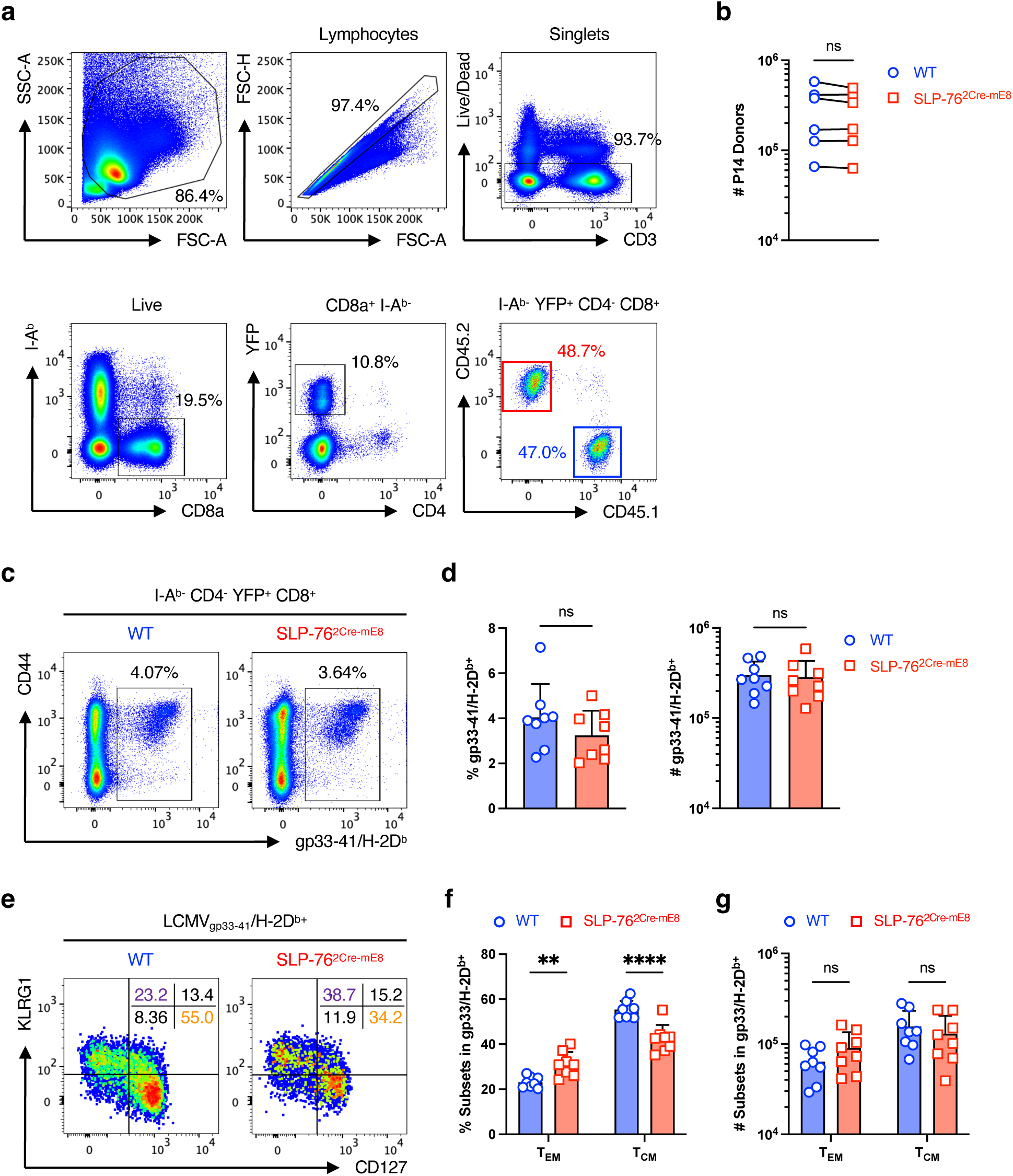
Gating strategy for analysis of donor P14 CD8^+^ T cells, and characterization of LCMV-reactive CD8^+^ T cells in LCMV_Arm_-infected WT and SLP-76^2Cre-mE8^ mice with endogenous TCR repertoire (corresponding to Figure 5). **a**,**b**, Gating strategy for analysis of co-transferred WT CD45.1 and SLP-76^2Cre-mE8^ CD45.2 P14 CD8^+^ T cells (**a**) and statistical analysis for the absolute numbers of the two donor P14 cells (**b**) in the spleens of (CD45.1 x CD45.2) F1 recipient mice at 28 days post-infection with LCMV_Arm_. **c**–**g**, WT and SLP-76^2Cre-mE8^ mice bearing endogenous TCR repertoire were infected with LCMV_Arm_. Thirty days post-infection, spleens were excised to analyze LCMV-reactive CD8^+^ T cells. **c**, Representative pseudocolor plot analysis of YFP^+^CD8^+^ T cells for gp33-41/H-2D^b^-tetramer binding and CD44 expression. **d**, Statistical analysis for the frequency (left) and absolute number (right) of gp33-41/H-2D^b^-tetramer^+^ CD8^+^ T cells (n=8). **e**, Representative pseudocolor plot analysis of gp33-41/H-2D^b^-tetramer^+^ CD8^+^ T cells for CD127 and KLRG1 expressions. The numbers indicate the frequency of cells in each quadrant. **f**,**g**, Statistical analysis of gp33-41/H-2D^b^-tetramer^+^ CD8^+^ T cells for the frequency (**f**) and absolute number (**g**) of T_EM_ (CD127^-^KLRG1^+^) and T_CM_ (CD127^+^KLRG1^-^) cells (n=8). Statistics was determined by paired t-test (**b**), two-sided Student’s t-test (**d**) and two-way ANOVA (**f**,**g**). ns; not significant, **; p<0.01, ****; p<0.0001.

**Extended Data Figure 6:**
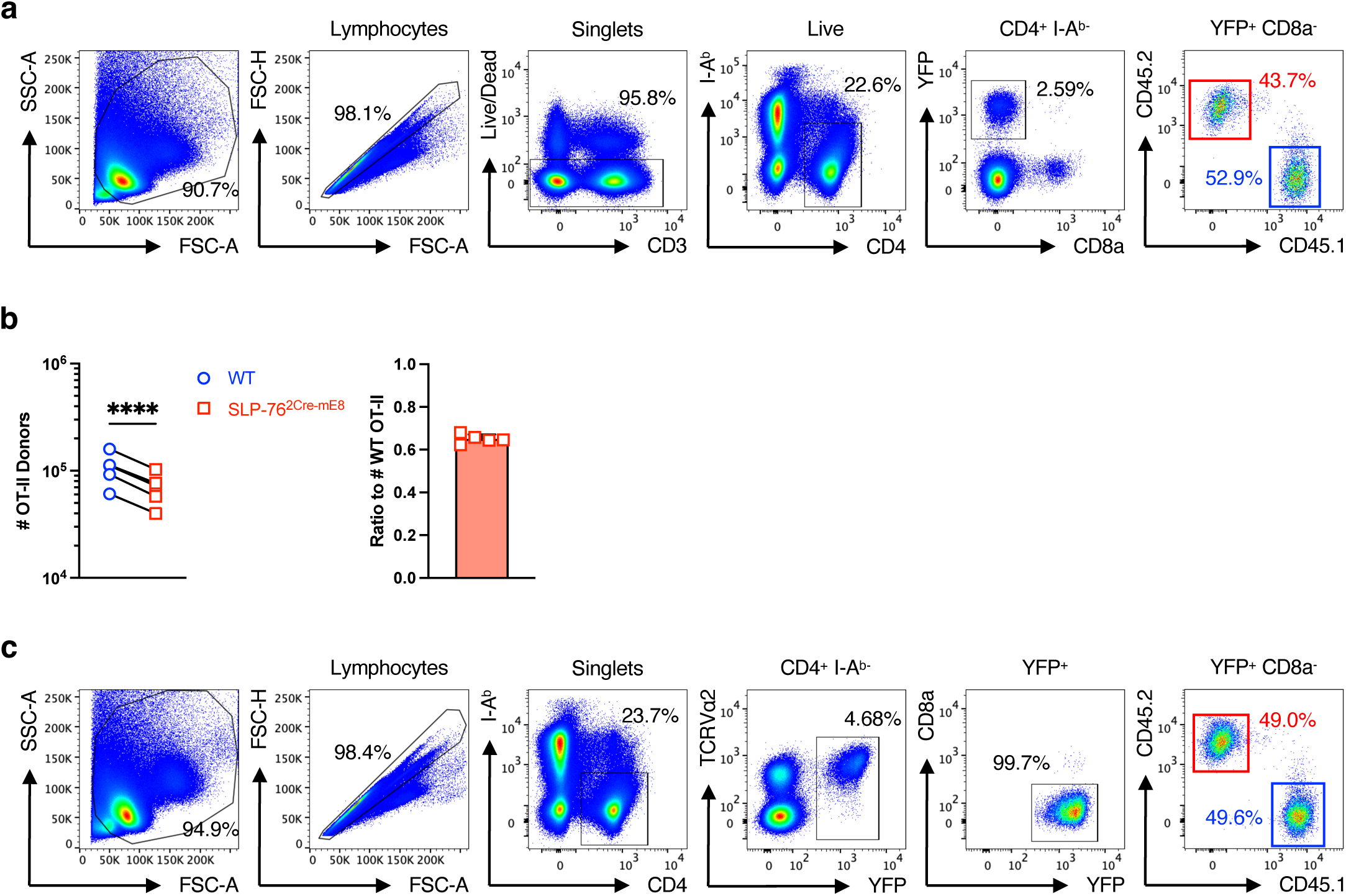
Gating strategies for analysis of donor OT-II CD4^+^ T cells for Tfh cells and for dead cells in a CXCR5^+^ population (corresponding to Figure 6). **a**, Gating strategy for analysis of co-transferred WT CD45.1 and SLP-76^2Cre-mE8^ CD45.2 OT-II CD4^+^ T cells in skin dLNs of (CD45.1 x CD45.2) F1 recipient mice at 10 days post-immunization with OVA in CFA. **b**, Statistical analysis for the absolute number of WT and SLP-76^2Cre-mE8^ OT-II donor cells (left) and the ratio of SLP-76^2Cre-mE8^ to WT OT-II donor cells (right) (n=5). **c**, Gating strategy for analysis of dead cells in CXCR5^-^ and CXCR5^+^ populations at 6 days post-immunization with OVA in CFA. Statistics was determined by paired t-test (**b**). ****; p<0.0001.

